# Homology-independent targeted integration *in vivo* restores *Cldn11* deficiency in mouse Sertoli cells and spermatogenesis

**DOI:** 10.64898/2025.12.23.696180

**Authors:** Tao Zhang, Anhao Guo, Hanben Wang, Yuan Chen, Lufan Li, Xin Wu

## Abstract

Defective testicular Sertoli cell (SC) function may be an underlying cause of male infertility/subfertility; however, *in vivo* gene therapy for nonobstructive azoospermia (NOA) caused by SCs has never been attempted. In this study, a CRISPR/Cas9-based homology-independent targeted integration (HITI) gene editing strategy was used to integrate an ∼3.7 kb DNA fragment containing a SC-specific enhancer/promoter and a region comprising three *Cldn11* exons into the SCs of *Cldn11*-deficient mice. After the viral plasmids carried by recombinant adeno-associated virus serotype 1 (rAAV1) were delivered to SCs via testicular tubular injection, the phenotypes of blood‒testis barrier loss, spermatogenesis blockage, and infertility in the mice were successfully rescued for 6 months, and first-generation offspring were produced using the sperm of the rescued mice. Our results suggest that despite the lack of proliferation and low division capacity of adult testicular SCs, the introduction of targeted strategies such as HITI could enable *in vivo* gene editing as a possible treatment for infertility caused by Sertoli cell defects.

## Introduction

Possible causes of infertility include not only deficiencies in the germ cells themselves but also functional and genetic abnormalities of the somatic cells in the gonadal microenvironment, which include Sertoli cells (SCs). SCs initiate testis formation during the embryonic period and are microenvironmental determinants of germ cell development. After birth, SCs can secrete neurotrophic factors, such as glial cell line-derived neurotrophic factor (GDNF), to promote the renewal of germline stem cells or lactate to support sperm maturation^1^. In addition, SCs provide physical structural support, for example, building the blood‒testis barrier (BTB), and mediate signaling between germ cells and the testicular microenvironment through pathways such as the CXCL12‒CXCR4 signaling axis^2^; both of these functions are crucial for germ cell homing and differentiation. There is growing evidence that dysfunction or genetic defects in SCs are potential causes of male infertility or subfertility in human patients^3^ and animal models, including in androgen insensitivity syndrome (AIS) caused by androgen receptor (AR) mutations^4^. Other gene mutations, such as *Hdac3*^5^, *Rac1*^6^, *Mex3b*^7^, and *Wt1*^8^, can also cause SC dysfunction.

Claudin-11 (CLDN11) is a key protein in epithelial cell junctions and was the first member of the claudin family to be knocked out and studied. The loss of CLDN11 leads to myelination dysfunction, hearing loss, and infertility in mice^9^. De novo mutation (stop codon variation) in the *CLDN11* gene leads to human hypomyelinating leukodystrophy^10^. In the testis, the integrity of tight junctions (TJs) depends on CLDN11 expression in SCs to ensure spermatogenesis^11^. Clinical evidence shows that the disruption of the BTB in patients with different degrees of spermatogenesis abnormalities is closely related to the aberrant expression of CLDN11^12^. In a previous study, transgenic mice were analyzed to identify cis-regulatory elements specific for *Cldn11* expression in SCs, and spermatogenesis was successfully rescued in *Cldn11*-deficient mice. The key cis-element is a 2 kb upstream DNA fragment starting from the *Cldn11* transcription start site^13^. Thus, it is possible to integrate exogenous DNA containing a complete regulatory element sequence into SCs through a gene editing strategy to achieve precise cis-expression of the CLDN11 protein in *Cldn11*-deficient mice.

CRISPR/Cas9-based gene editing technology using recombinant adeno-associated virus (rAAV) as a vector has been applied or has application prospects in the treatment of a variety of diseases. For example, the AAV5 vector, which has high tropism for photoreceptors, was used for gene editing to treat CEP290-associated retinal degeneration^14^, and AAV1-hOTOF-based dual-vector gene replacement therapy to correct mutations in the OTOF gene was tested in the first international clinical trial study of gene therapy for hereditary deafness^15^. In recent years, homology-independent targeted integration (HITI), a transgene knock-in strategy based on nonhomologous end joining (NHEJ), has been shown to efficiently knock in DNA sequence and increase the frequency of site-specific integration in both dividing and nondividing cells^16^. The HITI strategy was used to restore full-length dystrophin in the skeletal and cardiac muscle of Duchenne muscular dystrophy (DMD) model mice^17^, and HITI-mediated gene knock-in in mouse retinal photoreceptor cells was shown to improve X-linked juvenile retinoschisis caused by *Retinoschisin* 1 (*RS1*) mutations as well as retinitis pigmentosa (RP) caused by *Rhodopsin* (*RHO*) mutations^18, 19^. Nonetheless, there have been few reports of the use of gene editing strategies to address infertility or subfertility issues. Gene editing in germ cells may introduce DNA errors that can be passed to the next generation and thus raises distinct ethical issues^20^. Gene editing in testicular somatic cells is also difficult because somatic cells constitute only a small population of cells in the gonads, and some somatic cells stop dividing or proliferating after maturity^21^. In a recent study, transplantation of stem Leydig cells after prime editing-mediated *in vitro* correction of the *lhcgr* gene mutation was shown to rescue hereditary primary hypogonadism (HPH) in mice^22^. However, there have been no reports of *in vivo* gene editing for repair of gene deficiency or insufficient function of somatic cells in the niche of mammalian testis.

Here, we developed a DNA integration technique based on the HITI strategy and the CRISPR/Cas9 system in SCs *in vivo*, with plasmids carried by rAAV1. Through microinjection of rAAVs into the seminiferous tubules of *Cldn11*-deficient mouse testes, we achieved *in vivo* site-specific transgene integration in mouse SCs, restored CLDN11 expression, and rescued spermatogenesis and offspring production.

## Results

### Endogenous Cas9 expression in SCs by rAAV1 delivery and iCre induction

To achieve CRISPR/Cas9-mediated gene editing in SCs, we first established an inducible endogenous *Streptococcus pyogenes* Cas9 (SpCas9) expression system in SCs. We used *Rosa26^Cas9-tdTomato^* mice in which the transgenic sequence is knocked into the *Rosa26* locus and interrupted by a loxP-Stop (3 ×polyA signal)-loxP (LSL) cassette but can be driven by the CAG promoter under the induction of Cre recombinase (Figure 1A). We intended to isolate SCs from 18-day-old *Rosa26*-integrated *Cas9*-expressing transgenic mice (hereafter referred to as *Rosa26^Cas9-tdTomato^*^23^) because the SCs in the testis are mature at this stage, express hormone receptors and proliferate slowly (Figures S1A and S1B). Subsequently, we could obtain the overexpression of the Cre enzyme in SCs through an *iCre* plasmid carried by rAAV1 (rAAV1-*iCre,* a plasmid containing the *iCre* sequence driven by the EF1α promoter, 2222 bp), thereby achieving inducible expression of the Cas9 protein that can also be visualized by td-Tomato fluorescence (Figure 1A).

**Figure 1.**
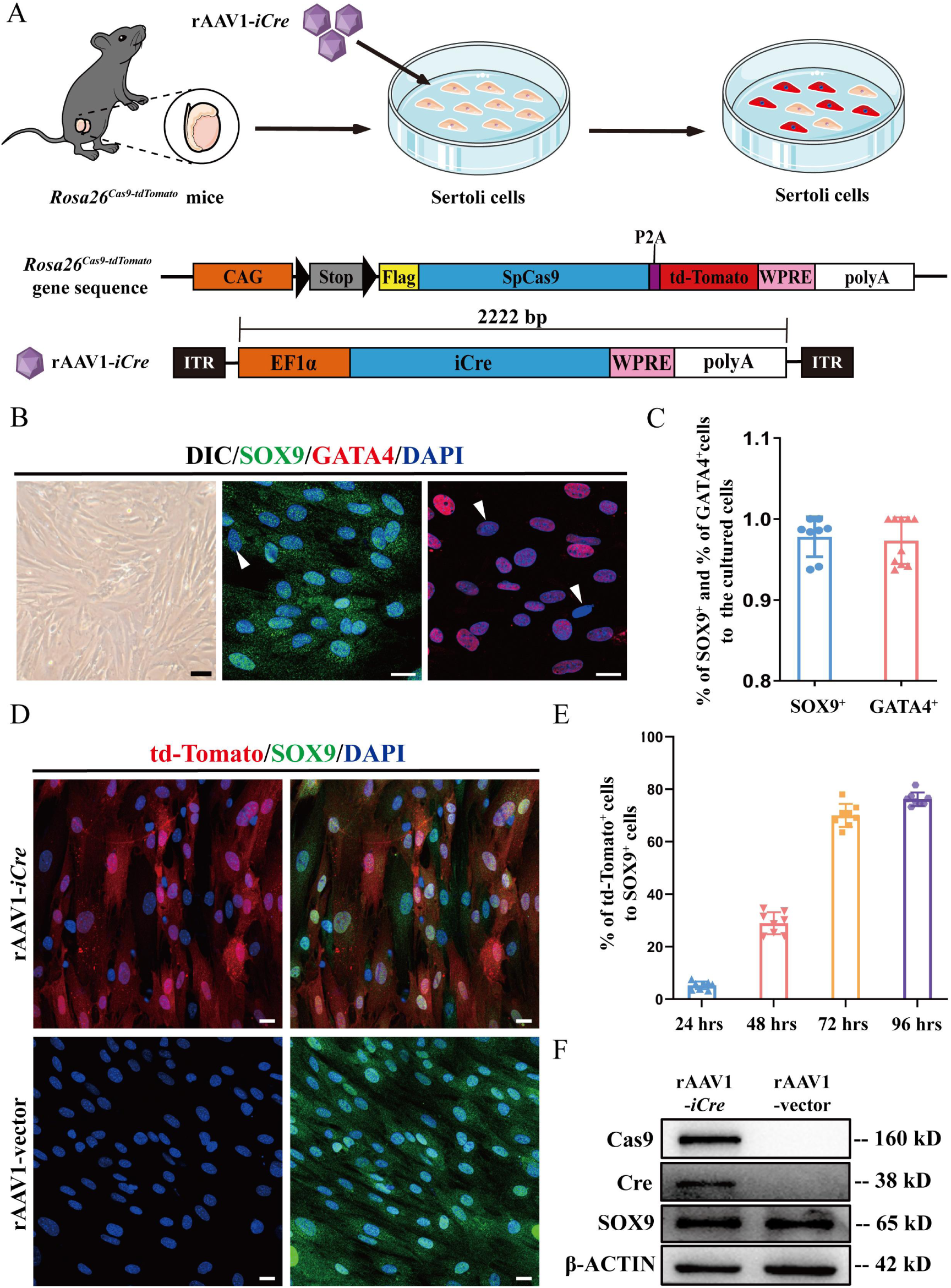
*In vitro* culture of primary Sertoli cells and rAAV1-mediated cell transfection. (A) Schematic diagram of rAAV1 transfection of SCs, the gene sequence of the *Rosa26^Cas9-tdTomato^* transgenic mice and the construction of the rAAV1-*iCre* plasmid. (B) DIC images (bright) and immunofluorescence staining for SOX9 (green) and GATA4 (red) in primary cultured *Rosa26^Cas9-tdTomato^*SCs. Arrows (white) indicate presumably negative cells. Scale bars: 50 μm in DIC and 20 μm in IF fields. (C) The percentage of isolated and cultured primary SCs was assessed by double staining for SOX9 and GATA4. Mann-Whitney U test was used for statistical analyses, U=36.500 and *Z*= -0.321. (D) Immunofluorescence colocalization of SOX9 (green) and td-Tomato (red) in primary cultured *Rosa26^Cas9-tdTomato^* SCs 72 h after rAAV1-*iCre* or rAAV1-vector transfection. Scale bar: 20 μm. (E) The percentage of td-Tomato^+^ cells to all SOX9^+^ SCs after rAAV1-*iCre* transfection; rAAV1-vector transfection was used as a control. Repeated measures ANOVA was used for statistical analyses, Geisser-Greenhouse correction was applied, GG=0.449 and *F*=688.838. (F) The protein expression of Cas9, Cre and SOX9 in *Rosa26^Cas9-tdTomato^*SCs detected via Western blotting at 72 h after rAAV1-*iCre* transfection; β-ACTIN was used as the protein loading control.

Examination of the rAAV1-*iCre* virus particles via transmission electron microscopy indicated that the viral samples contained 97.6 ± 0.4% full capsids and were suitable for transfection and subsequent experiments (Figures S1C and S1D). The AAV capsid protein was further verified by examining the ratio of the three capsid protein subunits (VP1:VP2:VP3), which was close to 1:1:10 (Figure S1E), as predicted^24^. Next, we transferred rAAV1-*iCre* into cultured mouse primary SCs and verified the cell purity through analysis of the marker proteins SOX9 and GATA4, which indicated that 97.8 ± 0.9% of cells were SOX9-positive and 97.3 ± 0.9% of cells were GATA4-positive (Figures 1B and 1C). A strong td-Tomato fluorescence signal appeared in the cultured SCs 72 h after transfection, and the proportion of positive cells increased from 5.3 ± 0.5% at 24 h to 76.2 ± 0.9% at 96 h after transfection (Figures 1D and 1E). Furthermore, Western blotting experiments verified the presence of Cre and Cas9 proteins in the transfected SCs compared with AAV-vector transfected SCs without Cre induction (Figure 1F).

### Testicular tubular injection enables efficient *in vivo* delivery of rAAV1-packaged recombinant DNA into SCs

To explore the best method for *in vivo* rAAV1 infection of SCs, we assessed the infection of testicular cells by observing the fluorescence intensity of the whole testis under a microscope after microinjection of rAAV1-*iCre* through the testicular rete (tubular injection), testicular interstitium (interstitial injection) or tail vein (tail vein injection) of *Rosa26^Cas9-tdTomato^* mice (Figure 2A). Fluorescence intensity analysis at multiple time points (5th, 10th, 20th, and 30th days) within one month after rAAV1-*iCre* injection indicated that both tubular injection and interstitial injection were effective for infecting testicular cells, whereas tail vein injection resulted in poor infection of the testis, although rAAV1-*iCre*-induced td-Tomato fluorescence enrichment was observed in the liver (Figures 2B and S3A). We further compared the number of td-Tomato-positive cells among the SOX9-positive SCs in mouse testes one month after rAAV1-*iCre* injection via three methods (Figure 2C) and found that tubular injection resulted in 75.7 ± 4.6% of SCs being efficiently infected, while the proportion of positive cells after interstitial injection was 42.1 ± 10.9%. In contrast, tail vein injection induced td-Tomato signal expression in SCs only rarely (Figure 2D). We also assessed the potential infection capacity of germ cells after rAAV1-*iCre* transfection by calculating the proportion of td-Tomato-positive signals in ZBTB16-positive spermatogonia (Figure 2C). The results were consistent with the pattern shown in SCs, where tubular injection resulted in 46.1 ± 8.6% positive cells, interstitial injection resulted in 14.7 ± 6.7% positive cells, and tail vein injection resulted in only rare infection of spermatogonia (Figure 2E).

**Figure 2.**
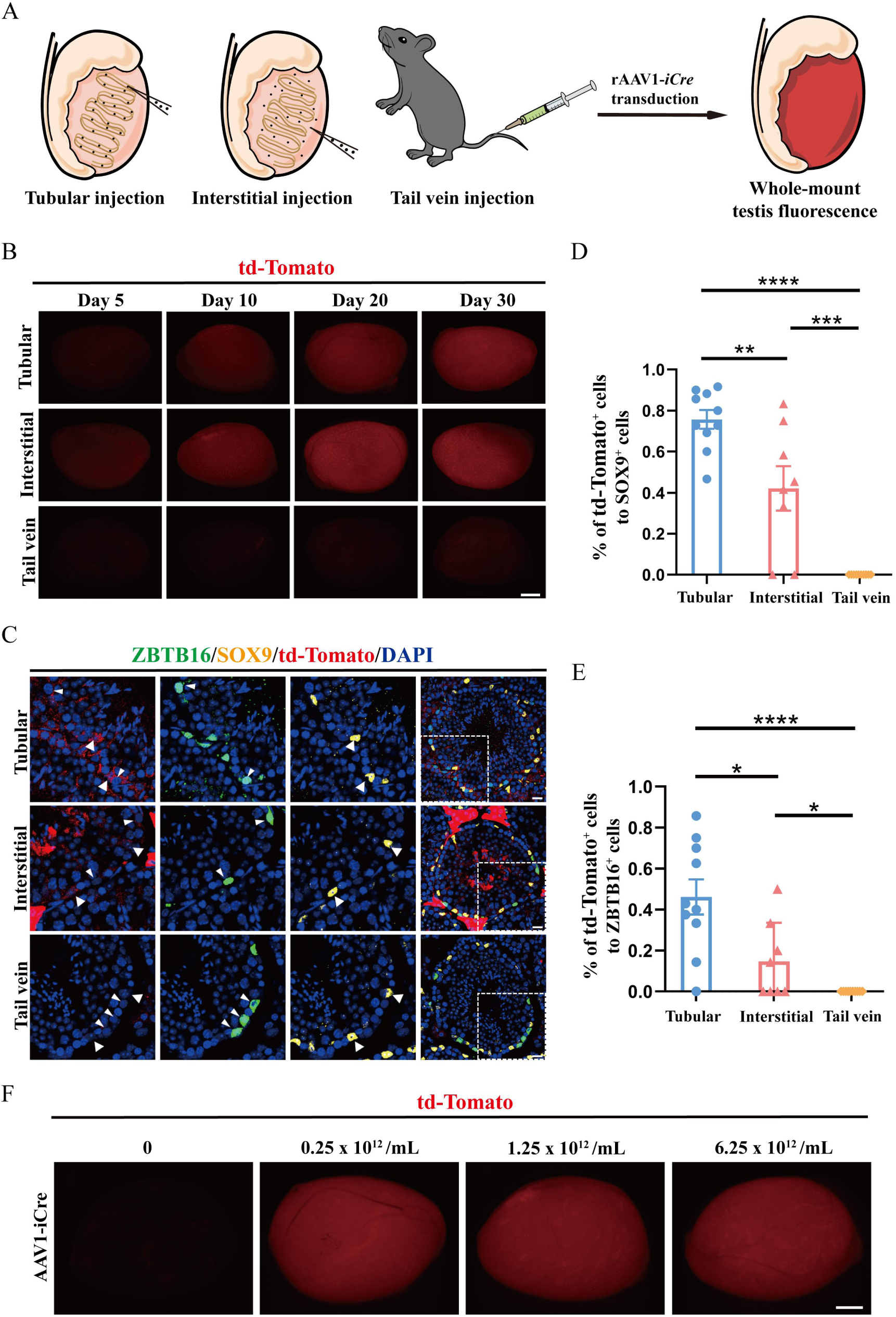
*In vivo* routes of viral delivery into mouse testes and titers applied for rAAV1 virus transduction. (A) Schematic diagram of rAAV1-*iCre* transduction into the testis via direct seminiferous tubules, interstitium, or tail vein. (B) Representative *Rosa26^Cas9-tdTomato^* mouse testes under a fluorescence microscope 40 days after rAAV1-*iCre* transduction. The intensity and localization of td-Tomato signals (red) are indicated in virus-transduced mouse testes through three different delivery routes. Scale bars: 1 mm. (C) Immunofluorescence colocalization of ZBTB16 (green), SOX9 (yellow) and td-Tomato (red) signals in mouse testes transduced with rAAV1-*iCre* through three virus delivery routes. ZBTB16^+^ cells are shown by small arrows, and SOX9^+^ cells are shown by larger arrows. Scale bars: 20 μm. (D) Comparison of the percentage of td-Tomato^+^ cells among SOX9^+^ cells in the testes of *Rosa26^Cas9-tdTomato^* mice. Kruskal-Wallis H test was used for statistical analyses, *H*=20.030. (E) Comparison of the percentage of td-Tomato^+^ cells among ZBTB16^+^ cells in the testes of *Rosa26^Cas9-tdTomato^* mice. Kruskal-Wallis H test was used for statistical analyses, *H*=16.261. (F) Intensity and localization of td-Tomato signals (red) in mouse testes after transduction with different titers of rAAV1-*iCre*. Scale bars: 1 mm.

Next, we evaluated the effects of the virus titer on the infection efficiency of testes. Three different titers of rAAV1-*iCre* (0.25 × 10^12^/mL, 1.25 ×10^12^/mL, and 6.25 ×10^12^/mL) were delivered by tubular injection, but we found no apparent difference in testis fluorescence intensity among them (Figure 2F). We selected a virus titer of 2.5 ×10^12^/mL for subsequent experiments as previously suggested^25^.

### The proximal 2 kb of the *Cldn11* promoter/enhancer region is sufficient to drive *in situ* expression in SCs

Next, we selected *Cldn11*-deficient mice (hereafter referred to as *Cldn11^KO^*) as a model to integrate exogenous DNA and rescue the lost function of SCs. *Cldn11^KO^* mice feature deletion of a 24.3 kb region containing three exons of *Cldn11* on chromosome 3 (Figure S2A), resulting in loss of CLDN11 expression, absence of testicular tight junctions, impaired spermatogenesis, and infertility^11^. To achieve *in situ* expression of CLDN11 in these mice, we utilized a regulatory region that has been previously shown to drive SC-specific expression as an endogenous promoter to drive expression of CLDN11 in mouse testes. This promoter region contained a 2 kilobase DNA fragment upstream (US 2 kb) of the *Cldn11* transcription start region^13^. We transduced rAAV1-packaged recombinant DNA containing the 2 kb promoter region and coding sequence consisting of three exons of *Cldn11* (a total of 3640 bp of *Cldn11*-donor sequence, rAAV1-*Cldn11*-donor, Figure 3B) into mouse embryonic fibroblast (MEF) cells (the infection of MEFs with rAAV1 was verified by EGFP signal in Figure S1F) and mouse primary SCs isolated from *Cldn11^KO^* mice. As above, rAAV1-*Cldn11*-donor particles were examined by transmission electron microscopy, and viruses containing 96.3 ± 0.3% full capsids were used for transfection, with a ratio of the three capsid protein subunits (VP1:VP2:VP3) close to 1:1:10 (Figures S1C‒S1E). Western blotting experiments revealed that the proximal 2 kb of the promoter/enhancer region was able to drive *in situ* expression of CLDN11 in SCs but not in MEF cells (Figure 3A). To validate whether this plasmid carried by rAAV1 could be effectively translated into functional CLDN11 protein *in situ* in mouse SCs, we analyzed mouse testes 40 days after tubular injection of the rAAV1-*Cldn11*-donor (Figure 3B). We found that the testicular weight of virus-infected mice was significantly increased, by approximately 55.9%, compared with that of their uninfected *Cldn11^KO^* littermates (40.7 ± 2.9 vs. 26.1 ± 0.3 mg; Figures 3C and 3D). Western blotting confirmed the presence of CLDN11, and immunofluorescence analysis of the basement membrane of the testicular seminiferous tubules further revealed that a tight junction barrier formed by CLDN11 could be detected in the testes of *Cldn11^KO^*mice 40 days after tubular injection (Figures 3E and 3F). However, the testis weights showed a decreasing trend with increased time after infection and approached the weights of *Cldn11^KO^* testes at 90 and 180 days after virus injection (31.7 ± 0.7 mg and 25.7 ± 1.8 mg, Figure 3D). Testicular histology examination confirmed that 40 days after infection, round and elongated spermatids were present in some seminiferous tubules of *Cldn11^KO^* mice, and spermatids were still found 90 days after infection (Figure 3G). However, as the time elapsed since the injection, the proportion of recovered seminiferous tubules continued to decrease, gradually returning to a degenerated morphology similar to *Cldn11^KO^* testes (66.3 ± 5.8% at 40 days, 52.2 ± 2.4% at 90 days, and 24.9 ± 3.9% at 180 days, Figures S5G and S5I). We used the spermatogonial marker ZBTB16, the spermatocyte marker γ-H2Ax, and the spermatid marker peanut agglutinin (PNA) to perform immunofluorescence staining of the transfected testes and confirmed that seminiferous epithelium morphology could be found in the testes of *Cldn11^KO^* mice infected with rAAV1-*Cldn11*-donor for 40 days, but not in that of their *Cldn11^KO^* littermates, which lacked spermatogenesis (Figures 3H, S4A and S4B). Compared with tubular injection of rAAV1-*Cldn11*-donor, the interstitial injection of the rAAV1-*Cldn11*-donor also produced spermatogenesis in the testis 40 days later but with rare PNA-positive round spermatids and no elongated spermatids in the testes, indicating that interstitial injection is ineffective as a delivery route to support the expression of CLDN11 in SCs or rescue spermatogenesis (Figures S5A and S5B). It has been reported that knockout of *Cldn11* results in increased CLDN5 expression in mouse testes^26^, we investigated CLDN5 expression and confirmed this phenomenon in the testes of *Cldn11^KO^* mice (the level of Sertoli cell marker GATA4 was used as a reference, Figure 3I and S5E). At the same time, we also observed that CLDN5 expression was downregulated in the testes of *Cldn11^KO^*mice 40 days after injection of the rAAV1-*Cldn11*-donor (Figures 3I and S5D).

**Figure 3.**
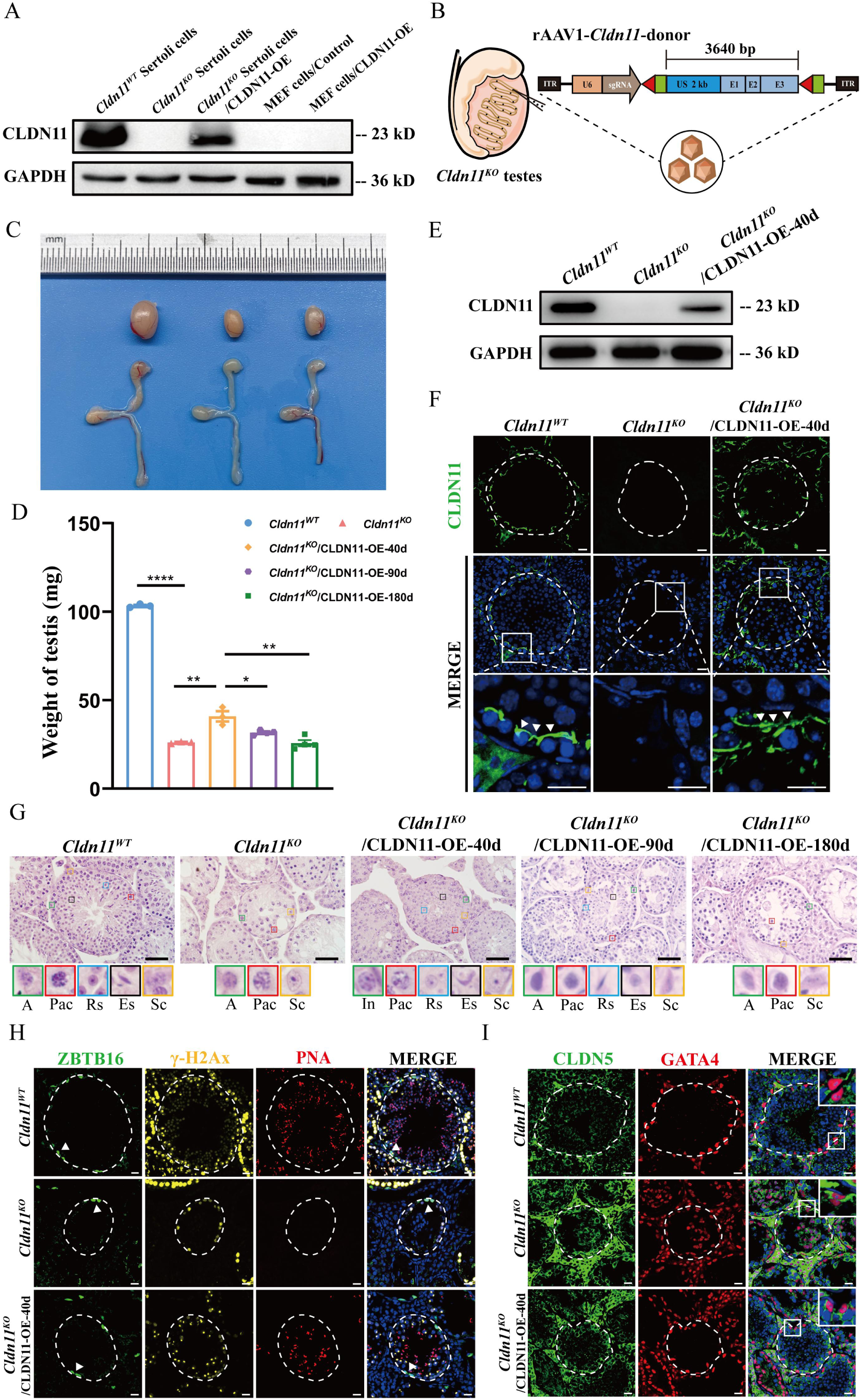
rAAV1-*Cldn11*-donor transduction transiently restores spermatogenesis in *Cldn11^KO^*mouse testes. (A) Induction of CLDN11 expression in *Cldn11^KO^* SCs and MEF cells transfected by rAAV1-*Cldn11*-donor were verified by Western blotting, compared to *Cldn11^WT^* and *Cldn11^KO^* SCs. GAPDH was used as the protein loading control. (B) Schematic graph of rAAV1-*Cldn11*-donor transduction to overexpress CLDN11 in *Cldn11^KO^* mouse testes via testicular tubular injection. (C) Representative morphological images of testes and epididymides from *Cldn11^WT^* mice, *Cldn11^KO^* mice and *Cldn11^KO^* mice 40 days after rAAV1*-Cldn11*-donor transduction. (D) Comparison of the testis weights of *Cldn11^WT^* mice*, Cldn11^KO^* mice and rAAV1-*Cldn11*-donor transduced *Cldn11^KO^* mice (n ≥ 3). One-way ANOVA was used for statistical analyses and *F*=417.865. (E) Induction of CLDN11 expression in *Cldn11^KO^* testes was verified by Western blotting, with comparison to *Cldn11^WT^* and *Cldn11^KO^* testes. GAPDH was used as the protein loading control. (F) Immunofluorescence of CLDN11 (green) in *Cldn11^WT^, Cldn11^KO^* and rAAV1-*Cldn11*-donor transduced *Cldn11^KO^* testes, shown by small arrows. Scale bars: 20 μm. (G) Testis histology revealed restored spermatogenesis in the testes of rAAV1-*Cldn11*-donor transduced *Cldn11^KO^* mice compared with those of *Cldn11^WT^* and *Cldn11^KO^* mice. The lower panels with colored frames show the types of cells found in the mouse testes. A, type A spermatogonia; In, intermediate spermatogonia; Pac, pachytene spermatocyte; Rs, round spermatid; Es, elongated spermatid; Sc, Sertoli cell. Scale bars: 50 μm. (H) Immunofluorescence of ZBTB16 (green), γ-H2Ax (yellow) and PNA (red) in *Cldn11^WT^, Cldn11^KO^* and rAAV1-*Cldn11*-donor transduced *Cldn11^KO^* testes. Scale bars: 20 μm. (I) Immunofluorescence of CLDN5 (green) and GATA4 (red) in *Cldn11^WT^, Cldn11^KO^* and rAAV1-*Cldn11*-donor transduced *Cldn11^KO^* testes. Scale bars: 20 μm.

Together, these findings indicated that the 2 kb enhancer and promoter sequences upstream of *Cldn11* transcription start region could efficiently drive *in situ* expression of CLDN11 in SCs, while the expression of CLDN11 seems to transiently support the restoration of spermatogenesis in *Cldn11^KO^* testes.

### Homology-independent targeted integration of recombinant *Cldn11* cassette into SCs via CRISPR/Cas9-mediated genome editing

Given the slowly dividing or non-proliferative nature of mature SCs, we investigated the possibility of using a CRISPR/Cas9-mediated HITI knock-in strategy to introduce exogenous DNA into the genome of *Cldn11*-defective SCs and achieve CLDN11 expression for long-term functional rescue of spermatogenesis. Through a breeding strategy (Figures S2A and S2B), we first constructed *Cldn11^KO^* mice that endogenously express Cas9 protein after induction with the Cre enzyme (rAAV1-*iCre*) for CRISPR/Cas9-mediated editing of SCs *in situ*; the resulting mice are hereafter referred to as *Cldn11^KO^-Rosa26^Cas9-tdTomato^* (Figure 4A). This approach enabled us to avoid nonspecific contamination and reduced integration efficiency caused by excessive loading of the rAAV1 packaging vector into SCs by eliminating the need to introduce a Cas9 expression system separately. Next, we selected a region approximately 8.5 to 9 kb upstream of the first exon of the *Cldn11* gene as the target for knock-in (Figure S6A). According to predictions from the NCBI gene database, gene integration in this region likely does not interrupt the expression of possible genes, and this region is located distant from the farthest region (∼5 kb upstream) of the endogenous regulatory elements of the *Cldn11* gene^13^. Next, in the plasmid of rAAV1-*Cldn11*-donor, we designed a *Cldn11*-donor sequence flanked by the same two sgRNA target sites and introduced a single guide RNA (sgRNA) sequence whose expression is driven by the human U6 promoter at the 5’ end of the *Cldn11*-donor sequence. In this case, the *Cldn11*-donor sequence could be integrated into the target site by *in situ* Cas9 cleavage under the guidance of sgRNA in SCs. Since the sgRNA target site sequences are in opposite orientations in the genomic DNA and rAAV1-*Cldn11*-donor, the sgRNA target sites are destroyed by forward integration (namely, donor integration generated by HITI, as shown in the red box, Figure 4B), and Cas9-based editing stops. Otherwise, the cleavage stage is re-entered until forward integration occurs (Figure 4B).

**Figure 4.**
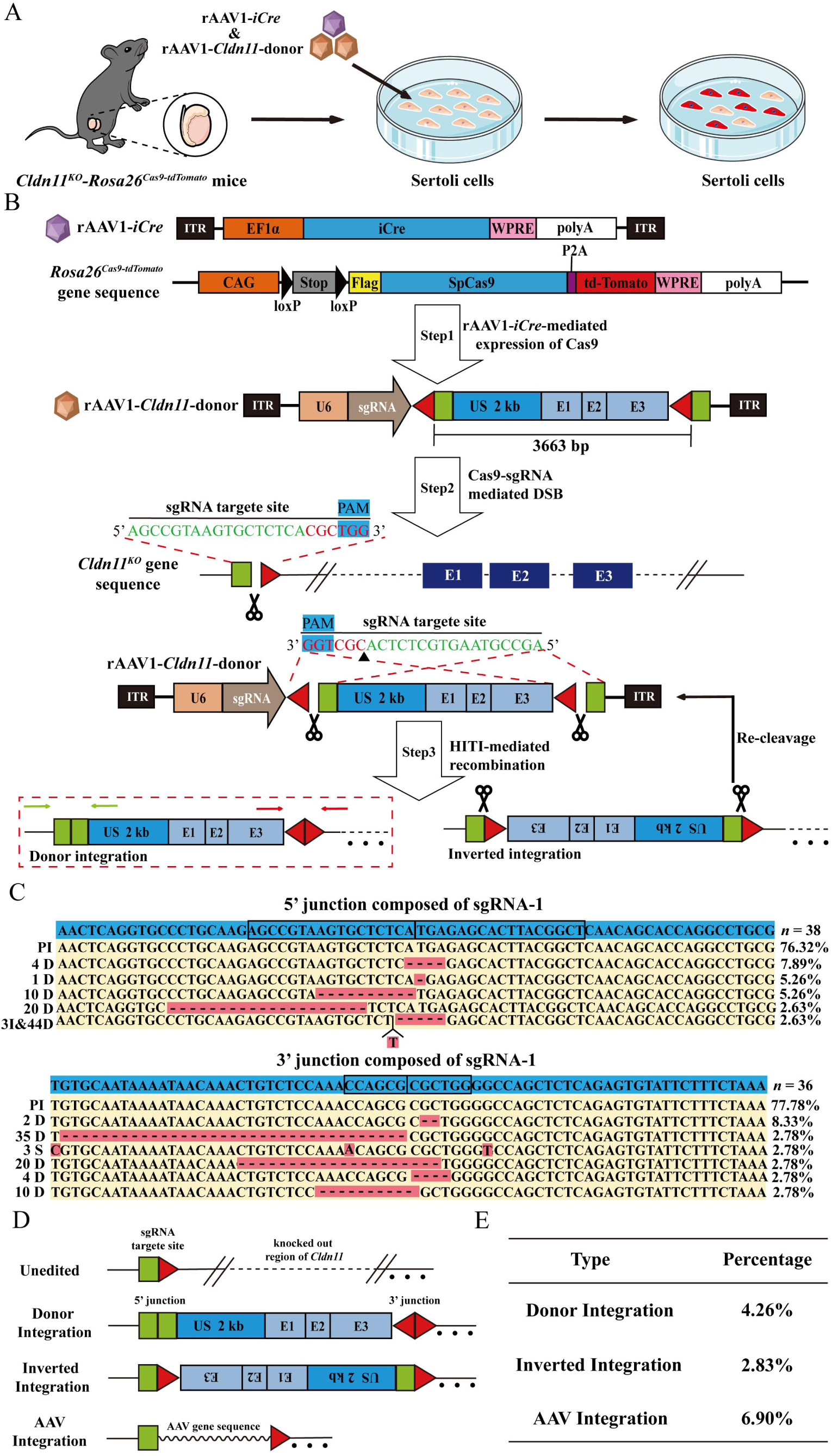
*Cldn11* integration at the target locus in mouse Sertoli cells using a HITI-mediated recombination strategy. (A) Schematic of rAAV1-*iCre* and rAAV1-*Cldn11*-donor virus transfection into cultured *Cldn11^KO^*-*Rosa26^Cas9-tdTomato^* SCs. (B) Schematic of Cas9-induced genomic integration by the HITI-mediated recombination strategy. US: upstream. (C) Sequencing of the 5’ and 3’ junctions generated by HITI-mediated recombination in *Cldn11^KO^-Rosa26^Cas9-tdTomato^* SCs. (D) Schematic of potential targeted genomic editing. (E) Primer -extension-mediated sequencing analysis of the genomic editing in *Cldn11^KO^*-*Rosa26^Cas9-tdTomato^* SCs transfected with rAAV1-*iCre* and rAAV1-*Cldn11-*donor.

To identify a suitable sgRNA to guide Cas9 for DNA cleavage, we uploaded the aforementioned 0.5 kb sequence as the target region for sgRNA design to an online tool (https://www.atum.bio/eCommerce/cas9/input) and obtained three candidate sgRNAs. Using the T7 endonuclease I cleavage assay for activity screening, we found that sgRNA-1 had the highest cleavage activity (16.5%, Figure S6B, the red arrows in the figure indicate two different lengths of cut DNA fragments), and using Sanger sequencing of genomic DNA from SCs (TM4 cell line) transfected with plasmids containing these three sgRNAs, we confirmed that sgRNA-1 initiated the most obvious DSB repair-mediated base loss (Figure S6C). Next, we transduced *Cldn11^KO^-Rosa26^Cas9-tdTomato^* SCs with rAAV1-*iCre* and rAAV1-*Cldn11*-donor containing sgRNA-1 (Figure 4A), sequenced the genomes of transfected SCs by Sanger sequencing and analyzed the 5’ and 3’ junction sites generated by HITI. The results revealed that the precise integration rates of the 5’ junction site and the 3’ junction site were 76.32% and 77.78%, respectively, confirming that the *Cldn11*-donor sequence was successfully integrated through the HITI strategy (Figure 4C). More importantly, most of the forward knock-ins did not result in indels, consistent with the results reported by Keiichiro Suzuki et al. in HEK293 cells^16^. Next, to comprehensively map all possible genome editing outcomes, we employed primer-extension-mediated sequencing (PEM-seq) as suggested^27^. PEM-seq showed that the editing efficiency of Cas9 in SCs at the sgRNA-1 target site was 29.8% (Figure S7A), which was higher than that predicted by the T7 endonuclease I cleavage assay (16.5%, Figure S6B). In addition, PEM-seq further revealed integration events including donor integration (4.26%), inverted integration (2.83%), and AAV integration (6.90%) (Figures 4D and 4E), and the overall donor integration efficiency in our study was more promising compared with that in another HITI editing study in skeletal and cardiac muscle^17^. Notably, both donor integration and inverted integration here could restore the expression of CLDN11 due to containing the SC-specific enhancer/promoter. In addition, to assess unwanted off-target effects caused by HITI editing, we applied a T7 endonuclease I cleavage assays of the four potential off-target sites of sgRNA-1 (up to 3 base mismatches), which revealed no significant off-target effects (red triangles indicate the predicted size of the cleaved DNA; Figures S6D and S6E). Consistently, we used PEM-seq to capture the off-target sites of sgRNA-1 and confirmed that no translocations of any off-target sites were observed (except for the translocation between the target site and the donor sequence, Figure S7B), and the major editing events were concentrated on the target site of sgRNA-1 (Figure S7C). As expected, we detected the successful protein expression of Cas9, Cre and CLDN11 in cultured *Cldn11^KO^-Rosa26^Cas9-tdTomato^* SCs (Figure S6F).

### Precise integration of recombinant *Cldn11* enables long-term rescue of spermatogenesis in *Cldn11^KO^* mice

Next, we explored whether integration of the *Cldn11*-donor sequence into the genome by the HITI strategy could enable SCs to express functional CLDN11 protein, thereby restoring spermatogenesis in the testes of *Cldn11^KO^-Rosa26^Cas9-tdTomato^* mice. We introduced both rAAV1-*iCre* and rAAV1-*Cldn11*-donor into the testes of *Cldn11^KO^-Rosa26^Cas9-tdTomato^* mice via tubular microinjection (Figure 5A) and, as expected, found that the testis weights were increased 40 days after virus injection compared with those of *Cldn11^KO^-Rosa26^Cas9-tdTomato^* littermates (40.3 ± 2.8 vs. 26.8 ± 0.3 mg, Figures 5B and 5C), which were similar to the testis weights 40 days after transient transfection with the rAAV1-*Cldn11*-donor alone. Consistent with this, we detected the protein expression of Cas9, Cre and CLDN11 in the testes 40 days after virus transfection (Figure 5D). However, in contrast to *Cldn11^KO^* testes with transient expression of *Cldn11*, the *Cldn11^KO^-Rosa26^Cas9-tdTomato^* testes with the *Cldn11* donor sequence integrated via HITI did not return to the littermate-like weight at later timepoints but rather maintained a weight roughly similar to that at 40 days at 90 and 180 days after microinjection (40.7 ± 3.7 and 44.0 ± 2.2 mg, respectively, Figure 5C). In line with the morphological data, we detected extensive, complete spermatogenesis and the presence of germ cells at various stages in mouse testes not only at 40 days after transfection (Figures 5E and S4C) but also at 90 days and 180 days after transfection (Figure 5G). Again, we performed immunofluorescence analysis of the basement membrane of the seminiferous tubules in *Cldn11^KO^-Rosa26^Cas9-tdTomato^* testes after virus transfection and observed a CLDN11-positive tight junction barrier (Figure 5F).

**Figure 5.**
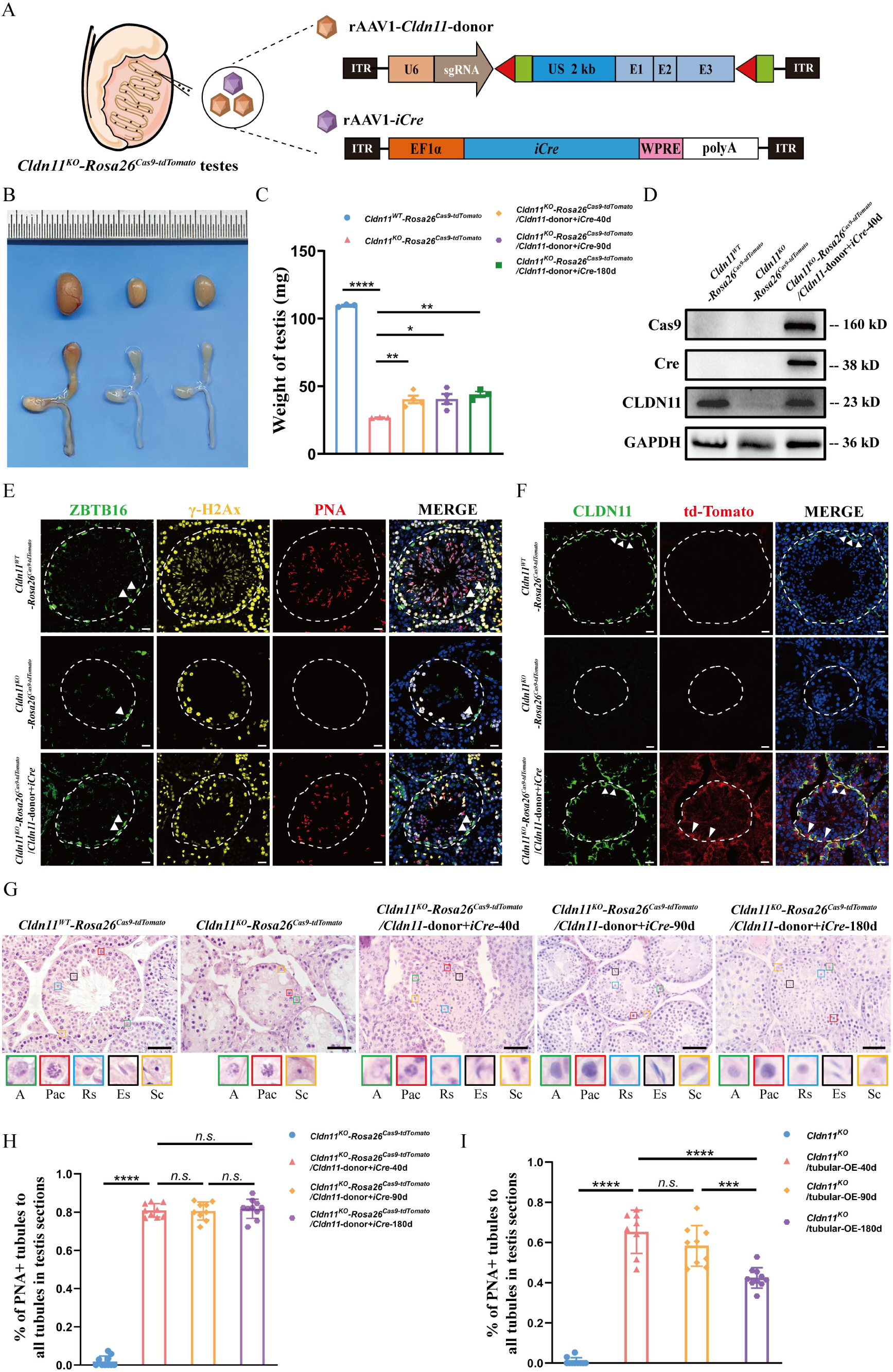
Genomic HITI of *Cldn11* rescues spermatogenesis in *Cldn11*-deficient mouse testes. (A) Schematics of testicular tubular injection of rAAV1-*iCre* and rAAV1-*Cldn11*-donor for validation of the HITI strategy-mediated *Cldn11* integration. (B) Representative morphological image of *Cldn11^KO^*-*Rosa26^Cas9-tdTomato^* testes and epididymides 40 days after tubular injection with rAAV1-*Cldn11-*donor and rAAV1-*iCre*, compared with those from *Cldn11^WT^*-*Rosa26^Cas9-tdTomato^* and *Cldn11^KO^*-*Rosa26^Cas9-tdTomato^* mice. (C) Comparison of the testis weights of *Cldn11^WT^*-*Rosa26^Cas9-tdTomato^* mice, *Cldn11^KO^*-*Rosa26^Cas9-tdTomato^* mice and *Cldn11^KO^*-*Rosa26^Cas9-tdTomato^* mice with HITI strategy-mediated *Cldn11* integration (n ≥ 3). One-way ANOVA was used for statistical analyses and *F*=137.444. (D) Western blotting verified Cas9, Cre and CLDN11 protein expression in *Cldn11^KO^*-*Rosa26^Cas9-tdTomato^* mouse testes with HITI strategy-mediated *Cldn11* integration compared with that in *Cldn11^WT^*-*Rosa26^Cas9-tdTomato^* and *Cldn11^KO^*-*Rosa26^Cas9-tdTomato^* mouse testes. GAPDH was used as the protein loading control. (E) Immunofluorescence of ZBTB16 (green), γ-H2Ax (yellow) and PNA (red) in *Cldn11^WT^*-*Rosa26^Cas9-tdTomato^, Cldn11^KO^*-*Rosa26^Cas9-tdTomato^* testes and *Cldn11^KO^*-*Rosa26^Cas9-tdTomato^* testes with HITI strategy-mediated *Cldn11* integration. Scale bars: 20 μm. (F) Immunofluorescence of CLDN11 (green) and td-Tomato (red) in *Cldn11^WT^*-*Rosa26^Cas9-tdTomato^, Cldn11^KO^*-*Rosa26^Cas9-tdTomato^* testes and *Cldn11^KO^*-*Rosa26^Cas9-tdTomato^* testes with HITI strategy-mediated *Cldn11* integration, shown by small arrows. Scale bars: 20 μm. (G) Extensive spermatogenesis in *Cldn11^KO^*-*Rosa26^Cas9-tdTomato^* mouse testes with HITI strategy-mediated *Cldn11* integration compared with that in *Cldn11^WT^*-*Rosa26^Cas9-tdTomato^* and *Cldn11^KO^*-*Rosa26^Cas9-tdTomato^* testes. The lower panels with colored frames show the types of cells found in the mouse testes. A, type A spermatogonia; Pac, pachytene spermatocyte; Rs, round spermatid; Es, elongated spermatid; Sc, Sertoli cell. Scale bars: 50 μm. (H) The percentage of PNA^+^ tubules in *Cldn11^KO^-Rosa26^Cas9-tdTomato^* testes transduced with rAAV1-*Cldn11*-donor and rAAV1-*iCre* for different lengths of time compared with that transduced with rAAV1-vector. Kruskal-Wallis H test was used for statistical analyses, *H*=21.870. (I) The percentage of PNA^+^ tubules in *Cldn11^KO^* testes transduced with rAAV1-*Cldn11*-donor for different lengths of time compared with that in *Cldn11^KO^*testes transduced with rAAV1-vector. Kruskal-Wallis H test was used for statistical analyses, *H*=30.468.

Most notably, in contrast to the loss of haploids in the seminiferous tubules observed at 180 days after transient transfection, PNA-positive haploids remained widely present in the seminiferous tubules of the *Cldn11^KO^-Rosa26^Cas9-tdTomato^* mouse testes 180 days after rAAV1-*Cldn11*-donor and rAAV1-*iCre* microinjection (Figure S5C). We calculated the proportion of PNA-positive tubules in *Cldn11^KO^-Rosa26^Cas9-tdTomato^* testes 180 days after virus microinjection and found that 81.8 ± 1.6% of the total tubules were still PNA-positive (Figure 5H). This finding was different from that of the testes with transient expression of the *Cldn11*-donor, in which only 42.4 ± 1.6% of the tubules were PNA-positive tubules after 180 days (Figure 5I). In addition, we performed CLDN11 immunofluorescence staining on the testes at 180 days after the two different treatments (Figures 3B and 5A) to evaluate the tight junction barrier. The results revealed that in *Cldn11^KO^* testes with transient expression of the *Cldn11*-donor, a considerable portion of tubules (54.2 ± 2.8%) presented abnormal CLDN11 fluorescence signal (as indicated by asterisks in Figure S5F), whereas only 19.1 ± 3.1% of *Cldn11^KO^-Rosa26^Cas9-tdTomato^* testes with *Cldn11*-donor integration exhibited this phenotype (Figures S5F and S5H); rather, most of the tubules retained complete seminiferous epithelial morphology (69.1 ± 5.2% at 40 days, 74.5 ± 3.1% at 90 days, and 66.2 ± 3.6% at 180 days; Figures S5G and S5I). Taken together, these data suggested that HITI strategy-mediated *Cldn11* integration enables long-term rescue of spermatogenesis in the testes of *Cldn11*-deficient mice.

### Sertoli cell-specific *in vivo* HITI allows assessment of editing accuracy

Since all infected cells in *Cldn11^KO^-Rosa26^Cas9-tdTomato^* testes may be induced by rAAV1-*iCre* to express the Cas9 protein, leading to DSBs and gene integration in the genome, we cannot rule out the potential risk of gene editing in germ cells. To address this problem, we generated *Cldn11^KO^-Amh^Cre^-Rosa26^Cas9-tdTomato^* mice through a breeding strategy (Figures S2A and S2C). These mice express the Cas9 protein in testicular SCs only under the induction of AMH-Cre (directed by SC-specific promoter elements of the gene encoding anti-Mullerian hormone) and produce td-Tomato in SCs (Figure 6A). Therefore, after the introduction of the *Cldn11*-donor, CRISPR/Cas9-mediated integration occurred only in SCs *in vivo* (Figure S8A). Similarly, we repeated the rAAV1-*Cldn11*-donor tubular microinjection experiments and found that the testis weight of *Cldn11^KO^-Amh^Cre^-Rosa26^Cas9-tdTomato^* recipient mice increased significantly at 40 days after virus injection (Figures S8B and S8C). CLDN11 protein was expressed in the testes (Figure S8D), germ cells of all stages (Figures S8E and S8F) and CLDN11-dependent tight junctions at the base of the seminiferous tubules were detected (Figure S8G), and PNA-positive haploids were present in the tubules (Figures S8F and S4D).

**Figure 6.**
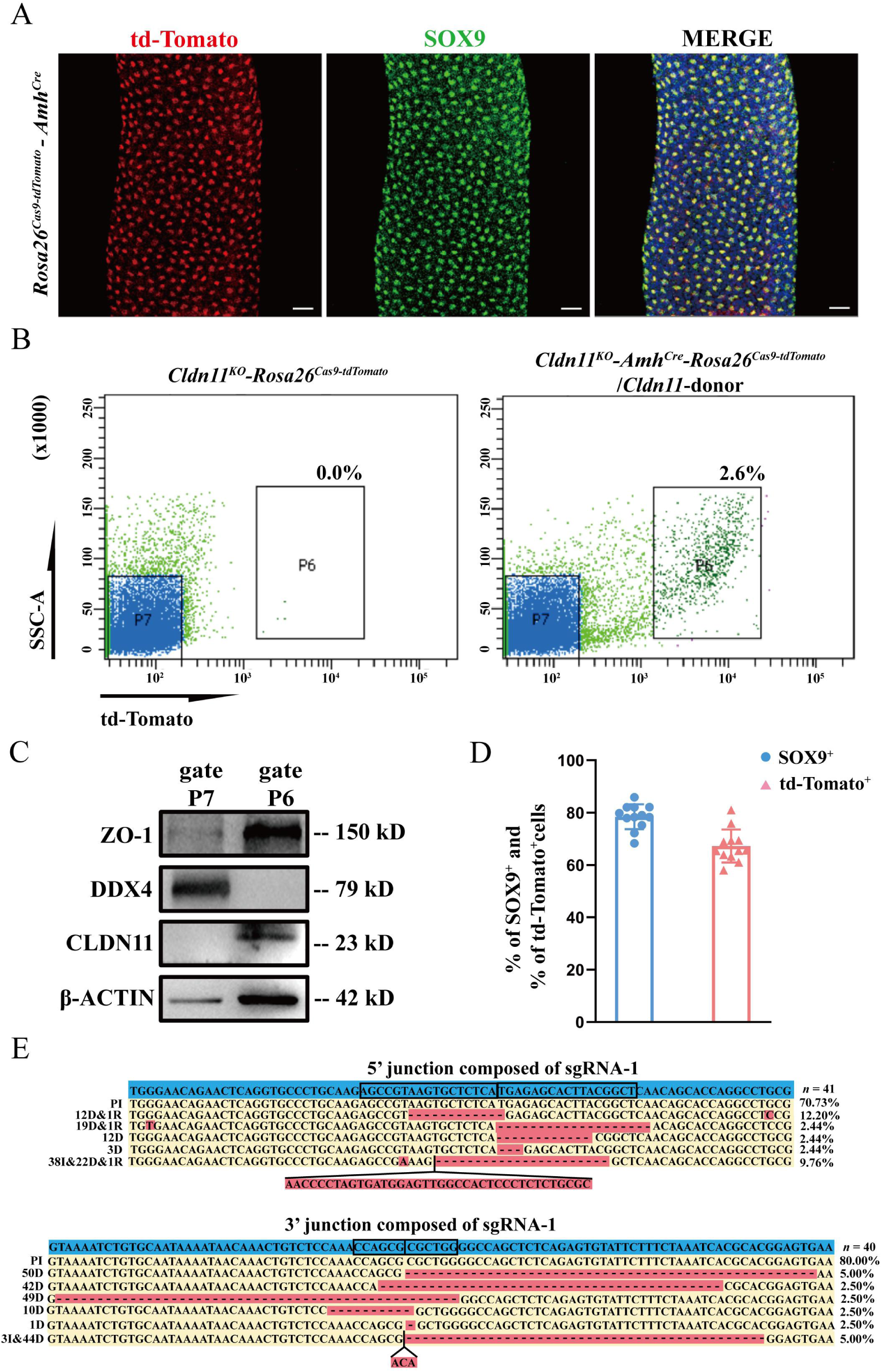
Assessment of *in vivo* SC-specific HITI and editing accuracy. (A) Whole-mount staining of SOX9 (green) in seminiferous tubules from *Rosa26^Cas9-tdTomato^-Amh^Cre^* mouse testes. td-Tomato (red) was autofluorescence from cells and colocalized with SOX9 (green) fluorescence. Scale bars: 50 µm. (B) Representative flow cytometry plots of td-Tomato expression in *Cldn11^KO^-Rosa26^Cas9-tdTomato^* testes *and Cldn11^KO^-Amh^Cre^-Rosa26^Cas9-tdTomato^* testes transduced with the rAAV1-*Cldn11*-donor. (C) Western blotting verified the protein expression of ZO-1, DDX4 and CLDN11 in cells sorted by flow cytometry. P6 and P7 represent td-Tomato-positive and td-Tomato-negative cell communities, respectively. The β-ACTIN level was used as the protein loading control. (D) Proportion of SOX9^+^ or td-Tomato^+^ cells among cells obtained via flow cytometry. Two independent samples T test was used for statistical analyses. (E) Sequencing results of the 5’ and 3’ junctions generated by HITI-mediated recombination in *Cldn11^KO^*-*Amh^Cre^*-*Rosa26^Cas9-tdTomato^* SCs.

Since genome cleavage and editing by CRISPR/Cas9 *in vivo* occurs only in SCs, it is possible to evaluate the accuracy of HITI at the DNA level in SCs after *in vivo* transfection. We obtained SCs expressing td-Tomato (td-Tomato fluorescence positive) from *Cldn11^KO^-Amh^Cre^-Rosa26^Cas9-tdTomato^* recipient mice by fluorescence-activated cell sorting (FACS) 40 days after virus injection (Figure 6B). After verifying the presence of CLDN11 in FACS-isolated cells by Western blotting (Figure 6C), we analyzed DNA extracted from SOX9-positive (78.4 ± 1.4%) or td-Tomato-positive (67.1 ± 1.7%) cells for the 5’ and 3’ junction sites generated by the HITI strategy (Figures 6D and S8H). The sequencing results again revealed that most of the forward knock-ins generated by *in vivo* HITI did not exhibit indels (precision integration rates at the 5’ junction and 3’ junction sites of 70.73% and 80.00%, respectively, Figure 6E).

### Sertoli cell-specific *in vivo* integration of *Cldn11* restores fertility and produces offspring

*Cldn11^KO^* mice are unable to mate naturally because of a hindlimb weakness phenotype^9^. Therefore, to assess whether the sperm produced in *Cldn11^KO^-Amh^Cre^-Rosa26^Cas9-tdTomato^* testes after gene therapy could support reproduction, intracytoplasmic sperm injection (ICSI) was performed via spermatozoa obtained by flow cytometry from male *Cldn11^KO^-Amh^Cre^-Rosa26^Cas9-tdTomato^* mice 40 days after rAAV1-*Cldn11*-donor injection, and oocytes were harvested from wild-type ICR female mice (Figures 7A and 7B). Twenty oocytes successfully progressed to the two-cell embryo stage *in vitro,* and these two-cell embryos (mixed with 15 wild-type two-cell embryos) were surgically transferred into the uteri of three pseudo-pregnant mice. A total of five newborn pups were obtained and applied to PCR-based genotyping (Figure 7C). The genotyping revealed that 3 of the 5 newborn pups carried mutated *Cldn11* alleles, and one carried the transgenic knock-in *Amh^Cre^* (Figure 7D). To test whether the rAAV1-*Cldn11*-donor was integrated into the genomes of the offspring, we designed specific primers for PCR to target the 5’- & 3’-ITR, U6 promoter and region containing the recombinant exons of *Cldn11* in the plasmid sequence (Figure 7E). We analyzed tail DNA samples from the three offspring, and the PCR results revealed no detectable vector sequences (Figure 7F). These findings indicated that the rAAV1-*Cldn11*-donor were not integrated into the genomes of the offspring.

**Figure 7.**
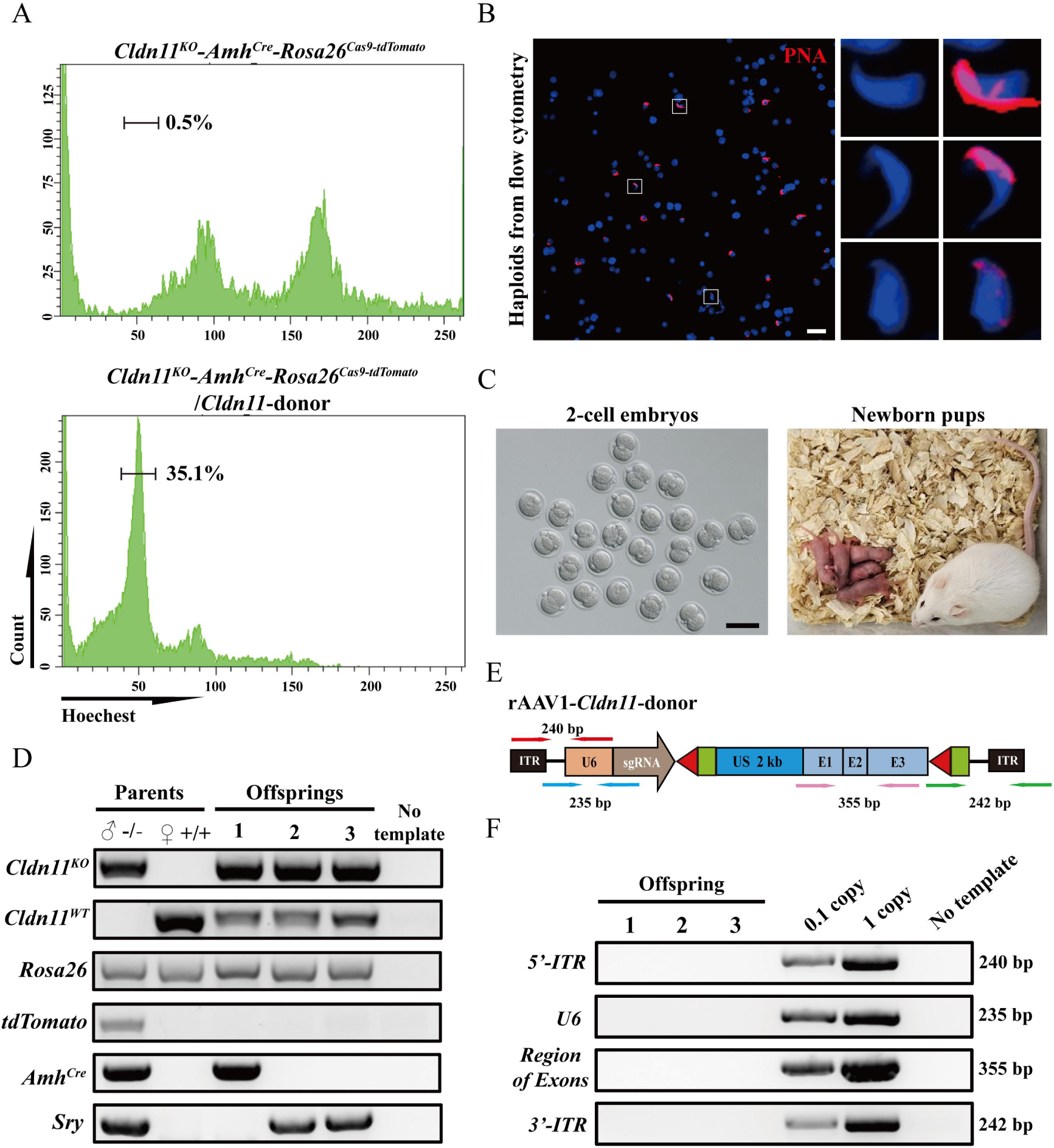
Generation of offspring after rescue of spermatogenesis via *in vivo* repair of *Cldn11* deficiency in infertile mice by HITI. (A) Representative flow cytometry plots of haploids in *Cldn11^KO^-Amh^Cre^-Rosa26^Cas9-tdTomato^* mouse testes transduced with rAAV1-*Cldn11*-donor compared with those in *Cldn11^KO^-Amh^Cre^-Rosa26^Cas9-tdTomato^* mouse testes. (B) Immunofluorescence of PNA (red) staining of haploids collected via flow cytometry from *Cldn11^KO^-Amh^Cre^-Rosa26^Cas9-tdTomato^* testes transduced with rAAV1-*Cldn11*-donor. Scale bars: 20 μm. (C) Representative images of 2-cell embryos and newborn pups. Scale bars: 50 μm. (D) Genotyping of the offspring derived from rAAV1-*Cldn11*-donor transfected *Cldn11^KO^-Amh^Cre^-Rosa26^Cas9-tdTomato^* mouse sperm. (E) Schematic diagram of primer design sites. The red, green, blue and pink arrows represent the orientations of the primers for the 5’-ITR, 3’-ITR, U6 promoter and the region of the recombinant exons of *Cldn11*, respectively. (F) PCR analysis of rAAV1-*Cldn11*-donor integration in the offspring genome. Viral particles representing 0.1 and 1 viral genome copies were used as controls.

## Discussion

In summary, we achieved genetic integration in SCs through a HITI-based strategy, restored CLDN11 expression after loss, and rescued the phenotype of testicular germ cell development arrest in *Cldn11*-deficient mice with spermatogenesis maintained for at least six months (see graphical abstract figure). Some previously reported mouse gene editing efforts to resolve fertility problems have focused on mouse germline stem cells^28–30^; however, human testicular germ cell gene editing for the generation of offspring is not feasible at present owing to the risk of transmission of unwanted genetic changes. Genetic defects in SCs are also the main cause of male fertility problems because they produce immature SCs that cannot support spermatogenesis^3^; dysfunction of the hypothalamus‒pituitary‒testis axis (HPT axis) and decreased androgen levels with testicular somatic cell ageing can also contribute to male infertility^31^. To our knowledge, this study is the first to demonstrate the use of cell-specific enhancers/promoters for *in vivo* gene therapy of male infertility in a mouse model, including non-obstructive azoospermia caused by genetic abnormalities in gonadal somatic cells.

Gene editing involves the use of nucleases, such as ZFNs (zinc finger nucleases), TALENs (transcription activator-like effector nucleases), or CRISPR/Cas9 (clustered regularly interspaced short palindromic repeats and the CRISPR-associated protein system), that target homologous sites in the genome and induce site-specific DNA double-strand breaks (DSBs)^32, 33^. Subsequently, cells repair DSBs by the NHEJ and/or HDR pathways. Although HDR-mediated repair can be used to introduce specific point mutations or insert desired sequences by initiating recombination of the target site with an exogenously provided DNA donor template, HDR can only occur during the S/G2 phase of the cell cycle, the repair efficiency is low, and this method cannot be used in nondividing cells^34^. NHEJ-mediated repair is active throughout the cell cycle in a wide range of somatic cells, but this repair is imprecise and can produce insertions and/or deletions (indels) of varying lengths at DSB sites; it is also homology-independent, and the direction of insertion of the exogenous DNA donor template cannot be controlled^33^. Therefore, a more efficient NHEJ-based strategy is needed to explore the feasibility of gene editing in extremely low-proliferation or nondividing cells such as SCs after male puberty.

In recent years, a new CRISPR-NHEJ-mediated gene knock-in method called homology-independent targeted integration (HITI) was developed by Keiichiro Suzuki et al. and was shown to be effective in both dividing and nondividing cells *in vivo* and *in vitro*^16^. Since then, multiple studies have demonstrated that the HITI strategy can be used to knock DNA into the genome of different tissue cells to treat various diseases. For example, *in vivo* HITI-mediated gene editing via intravenous injection of AAV9 vectors was used to rescue the pathogenic mutation of the X-linked ABCD1 gene in adrenoleukodystrophy (ALD) model mice, which showed that HITI is a more universal method than others because it is effective regardless of the mutation type or site^35^. Similarly, CRISPR/Cas9-mediated AAV-HITI targeting retinal photoreceptors improved retinal morphology and function in mice with autosomal dominant retinitis pigmentosa; AAV-HITI achieved stable transgene expression and phenotypic improvement in the liver of a mouse model of severe lysosomal storage disease mucopolysaccharidosis type VI^19^; CRISPR/Cas9-mediated HITI achieved permanent integration of a highly specific active factor IX variant at the albumin gene locus in a rat model of hemophilia B and improved hemophilia^36^; and AAV6-delivered, CRISPR/Cas9-mediated, and HITI-based transgene integration in the *ITGB2* gene was shown to effectively target human CD34^+^ hematopoietic stem and progenitor cells (HSPCs)^37^. In this study, we used rAAV1 delivery and CRISPR/Cas9-HITI-based strategies to integrate 3663 bp recombinant *Cldn11* donor gene sequences into SCs and used the 2 kb sequence upstream of the endogenous promoter to successfully achieve long-term expression of CLDN11 protein in SCs. This approach restored spermatogenesis in *Cldn11^KO^* mice over an observation period of at least 6 months. In addition to the HITI strategy, other novel gene integration strategies are gradually being developed and applied. For example, the primer editing-assisted site-specific integrase gene editing technology (PASSIGE) developed by David R. Liu’s laboratory combines primer editing with the ability of recombinase to accurately integrate large sequences of more than 10 kilobases^38^, whereas another strategy called LOCK (Long dsDNA with 3’-Overhangs mediated CRISPR Knock-in) can also efficiently target and integrate kilobase-sized DNA fragments^39^. The development of these strategies provides hope for achieving efficient gene integration and treating most genetic diseases.

Gene editing technology combined with recombinant adeno-associated virus is a promising therapeutic option because AAV is nonpathogenic and can infect both dividing and nondividing cells without significant cytotoxicity or the introduction of genomic insertion mutations^40, 41^. In the present study, we found 6.90% AAV genome fragment integration through PEM-seq (Figures 4D and 4E), and the side effects of this integration may need further evaluation. On the other hand, gene transduction into SCs can be achieved by microinjection of AAV into the seminiferous tubules; AAV1 and AAV9 can penetrate not only the testicular basement membrane but also the BTB and efficiently infect SCs^25^. Our SC transfection efficiencies by tubular injection and interstitial injection with rAAV1 were approximately 75.7% and 42.1%, respectively (Figure 2D), which is consistent with the efficiency previously reported in the literature^25^. In contrast, we found that after tail vein injection, rAAV1 was expressed mainly in the liver, which may be due to its weaker vascular penetration ability^42^. In addition, since different AAV serotypes have unique cell-targeting properties^43, 44^, further optimization of AAV serotypes may be needed.

When targeting gene therapy for specific diseases, it is highly advantageous to restrict gene expression to the tissue of interest. For example, human cardiac troponin T (*TNNT2*) promoter drives the expression of *MYBPC3* to treat hypertrophic cardiomyopathy (HCM)^45^. In this study, we utilized a 2 kb upstream sequence of the *Cldn11* transcription start site as a promoter-enhancer element to drive *Cldn11* expression. This made it possible to mimic the spatiotemporal expression pattern of the gene itself in Sertoli cells. As shown in Figure 3A, the unique promoter element of *Cldn11* ensures its specific expression in Sertoli cells, avoiding the potential immune response caused by ectopic protein expression. Similar experiments have been demonstrated in gene therapy for muscular dystrophy^46^. Furthermore, this approach maintains physiological levels of protein expression, preventing excess protein production that could cause cytotoxicity. By eliminating strong promoters like CMV, the risk of unintended tumor formation during gene integration (for example, if a strong promoter is accidentally integrated near an oncogene) can be avoided. In addition, some strong viral promoters are gradually silenced by the cell’s own defense mechanisms, whereas gene-intrinsic promoters can effectively circumvent this problem. Despite this, our strategy in this study still has certain limitations. First, the *Cldn11* donor sequence packaged in the rAAV1 plasmid in this study was approximately 4.4 kb (5’-ITR to 3’-ITR region), which is close to the upper limit of AAV packaging. When the sequence length exceeds 5 kb, the transfection efficiency of AAV decreases sharply^47, 48^. We also found that the expression level of CLDN11 in the testes of *Cldn11^KO^* mice still did not approach wild-type levels. These issues prompted us to further explore ways to shorten the promoter sequence while enhancing protein expression, such as using enhancer remodeling strategies. Besides, a dual AAV strategy (splitting the larger editing protein into two inactive fragments and packaging them into two AAVs for delivery) may be used; additionally, more compact gene editors for single AAV delivery, such as Cas12 variants and mandatory mobile element-directed activity (OMEGA) nucleases, have been developed^49^. Other delivery materials, such as nonviral lipid nanoparticles (LNPs), have strong mRNA encapsulation capabilities, although LNP targeting of most non-liver tissues remains a challenge^50^. In recent years, engineered virus-like particles (eVLPs) that combine the advantages of viral and nonviral vectors have been shown to achieve the effective delivery of gene editing tools to different organs of mammals^51, 52^; engineered PNMA2 (ePNMA2) particles or proteolipid vectors (PLVs) are also potential delivery vectors^53, 54^.

It should be noted that although this study successfully integrated the *Cldn11*-donor gene sequence using AAV-mediated HITI technology and restored the expression of CLDN11 in *Cldn11^KO^* SCs, our deep sequencing results showed that the efficiency of genome correction needs to be improved (Figure 4E). Improving the efficiency of HITI could help this long-fragment gene integration therapy strategy become more suitable for clinical applications. In recent years, several teams have successively developed a variety of Cas9 variants with low risks of off-target activity and improved on-target gene editing efficiency, including FrCas9, CoCas9 and iGeoCas9^55–57^. With the development of more gene editors and further improvements in editing efficiency, the HITI-mediated transgenic knock-in strategy can be expected to have wider applications in the future.

## Methods and materials

### Animals

*Cldn11^KO^* mice (NO. T032226) were purchased from GemPharmatech (Nanjing, China). According to the structure of the *Cldn11* gene, exon1-exon3 of the *Cldn11*-201 (ENSMUST00000046174.8) transcript, which contains all of the *Cldn11* coding sequence, was recommended as the knockout region, as knockout of this region was predicted to result in complete disruption of protein function. The results of gene knockout were as follows: 5’-ctaaggacaatccttgttcttactgccttt-24341 bp-aagcatacctcaaacaaggtgcggtcagta-3’. The *Rosa26^Cas9-tdTomato^* (NO. T002249) was purchased from GemPharmatech (Nanjing, China). A Cas9 expression element driven by the CAG promoter was inserted into the *Rosa26* site, and the STOP elements (3×polyA signal) flanked by *loxP* between the CAG promoter and the *Cas9* expression element were inserted to block Cas9 expression. In addition, the red fluorescence marker td-Tomato was incorporated downstream of the *Cas9* expression element to facilitate the detection of Cas9 expression. When Cre recombinase is present, cells express the Cas9 protein and td-Tomato red fluorescence. The *Amh^Cre^* mice were gifted by Xuejiang Guo (State Key Laboratory of Reproductive Medicine and Offspring Health, Nanjing Medical University, Nanjing, China). The PCR primers used for *Cldn11, Rosa26^Cas9-tdTomato^ and Amh^Cre^* genotyping are listed in Table S1, and the results of the genotyping are shown in Figure S2A. All the mice with a C57BL/6 genetic background used in this study were raised in a germ-free environment at the Experimental Animal Center of Nanjing Medical University. The mice were raised under a 12/12-h light‒dark cycle and provided sterile water and food as needed. The use of mice in this study was approved by the Institutional Animal Care and Use Committee of Nanjing Medical University (IACUC).

### Plasmid construction and rAAV production

To construct the sgRNA plasmids, the CRISPR-v2 plasmid (VT8107, YOUBIO) was digested with BsmB I-v2 (R0739, NEB), and oligos containing the target sequences were annealed and cloned into the CRISPR-v2 vector with T4 ligase (M0202, NEB). The sgRNAs used in this study are shown in Figure S6C. To construct pAAV-U6-sgRNA1-*Cldn11* plasmids, we used the CDS region of the *Cldn11* gene as the body of the alternative gene, connecting the 2000 bp promoter region upstream, to form the alternative *Cldn11* gene cassette. We subsequently used the sgRNA-1 recognition site flip sequence as the directional arm for HITI integration and attached it to both sides of the alternative *Cldn11* cassette to prepare pAAV-U6-sgRNA1-*Cldn11* using the pX552 (LM8197, LMAI Bio) plasmid backbone. To construct pAAV-EF1α-*iCre* plasmids, the EF1α promoter sequence and *iCre* sequence were amplified via PCR using the pLV-EF1α-MCS-IRES-Bsd (LM8179, LMAI Bio) and pCAG-Cre (LM9007, LMAI Bio) plasmids as templates and then ligated to pAAV-MCS (LM1520, LMAI Bio) digested with Mlu I-HF (R3198, NEB) and BamH I-HF (R0136, NEB) as the pAAV-EF1α-*iCre* backbone. Specifically, during the production of recombinant adeno-associated virus (rAAV), a helper plasmid containing adenoviral elements (pHelper), a plasmid containing the Rep and Cap genes, and a plasmid containing the ITRs were transfected into HEK293T cells. Viral replication and release were completed during the cell culture process. When sufficient viral particles were produced in the cell culture medium, the cells were lysed and the supernatant was collected. Finally, the viral particles were purified by ultracentrifugation. The rAAV titer was determined by quantitative qPCR to detect the copy number of exogenous DNA in the viral genome.

### SgRNA screening and validation of cleavage activity at on-target and off-target sites via a T7 endonuclease I cleavage assay

Three sgRNAs with minimal predicted off-target effects were designed by an online tool (https://www.atum.bio/eCommerce/cas9/input) to target the 500 bp region ∼8.5 kb upstream of *Cldn11* Exon-1, and the SpCas9 activity guided by the three sgRNAs in SCs was evaluated by using the T7EI assay. An online search tool (https://chopchop.cbu.uib.no/) was used to search the mouse genome (GRCh39) for potential SpCas9 off-target sites (PAM: 5’-NGG-3’) with the sgRNA-1 used in this study (5’-AGCCGTAAGTGCTCTCACGC-3’) as a query sequence, allowing up to 3 base pair mismatches between the sgRNA and candidate target sites. Only four potential off-target sites with three mismatches were identified (shown in Figure S6D), and no sites with 1 or 2 mismatches were identified. SCs were transfected with 125 ng of the *U6-sgRNA CRISPR-V2* plasmid using liposomes (40802ES03, Yeasen) according to the manufacturer’s protocol. The cells were incubated for 72 h, and genomic DNA was isolated with a DNA mini kit (51304, QIAGEN). Indels generated by the three sgRNAs at the four potential off-target sites were identified via PCR of the region of interest (primers provided in Table S1) via Super-Fidelity DNA Polymerase (P505, Vazyme), followed by purification (DP204-02, TIANGEN) and quantification of the DNA, incubation with the T7 endonuclease (EN303, Vazyme) and electrophoresis on a 1.5% agarose gel. Then, percent editing was calculated by measuring the intensities of the unedited and edited amplicon bands via ImageJ (1.8.0_345). These values were used to calculate the percentage of edited amplicon bands ([edited band 1 + edited band 2]/[total band intensities]) and then used to calculate percent editing by the following formula: (1 [square root (1 percentage of edited amplicon bands)] × 100)^17^.

### *In vitro* culture and rAAV1 transfection of mouse primary SCs and MEFs

Testes from 14–18 d mice were kept in 1 × PBS, and the tunica albuginea was stripped under a stereoscope. The mass of the seminiferous tubules was stretched and incubated in 5 mL of DMEM/F12 medium (11320-033, Gibco) containing 1 mg/mL collagenase IV (C5138, Sigma‒Aldrich) and 1 mg/mL hyaluronidase (H3884, Sigma‒Aldrich) for 5 min at 37°C with gentle shaking. The digested mixture was left at room temperature for 1 min to obtain separate seminiferous tubules. Then, the seminiferous tubules were washed in HBSS 3 times and incubated in 3 mL of 0.25% trypsin-EDTA (25200-056, Gibco) containing 1.5 mg/mL deoxyribonuclease I (DN25, Sigma‒Aldrich) for 5 min at 37°C with gentle shaking, and the reaction was terminated with fetal bovine serum. The cell suspension was filtered with a 40 μm cell strainer and centrifuged at 600 ×g for 5 min at 4°C, and the supernatant was discarded. The cells were transferred into DMEM/F12 medium (11320-033, Gibco) containing 10% FBS (A5670-701, Gibco), 1% GlutaMax (35050-061, Gibco) and 1% penicillin‒streptomycin solution (15140-122, Gibco) at 37°C with 5% CO2. After 24 h of culture, the cells were treated with 37°C hypotonic solution (20 mM Tris, pH = 7.4) for 2 min to remove residual germ cells and then incubated in fresh medium. MEFs were cultured in DMEM medium (11965-084, Gibco) containing 10% FBS (A5670-701, Gibco) and 1% penicillin‒streptomycin solution (15140-122, Gibco) at 37°C with 5% CO2. For rAAV1-*iCre* transfection of *Rosa26^Cas9-tdTomato^* primary SCs, viruses were added to the plates at an MOI of 5 × 10^4^ total vectors per cell. To validate the *in vitro* integration of *Cldn11* by the HITI strategy, rAAV1-*iCre* and rAAV1-*Cldn11*-donor were combined at a ratio of 1:2 and added to the plates at an MOI of 2 × 10^5^ total vectors per cell. The cells were cultured for several days and collected for analysis of genomic DNA, Western blotting and immunofluorescence.

### Analysis of 5’ and 3’ integration generated by the HITI strategy

The cultured primary SCs (14 days after rAAV1-*iCre* and rAAV1-*Cldn11*-donor transfection) or SCs collected by flow cytometric sorting were subjected to DNA extraction. The regions of interest were amplified via PCR using the primers listed in Table S1. The DNA fragments were recovered and purified with a gel purification kit (DP209, TIANGEN) and then ligated to *PClone007* (TSV-007S, TSINGKE). Then, we transformed the plasmid into *Trelief-5α* chemically competent cells (TSC-C01, TSINGKE), cultured the transformed cells on Luria–Bertani plates overnight in a 37°C incubator, and sent the plates with bacterial clones to TSINGKE for Sanger sequencing.

### Microinjection of recombinant adeno-associated virus particles

Male mice at 5–6 weeks were anesthetized by the intraperitoneal injection of 0.02 mL of 1.25% Avertin (Sigma‒Aldrich, T48402) per mg body weight, followed by the injection of 200 μL of a mixture of rAAV1 and HBSS by the tail vein. For testicular tubular microinjection and interstitial microinjection, virus particles were introduced into the seminiferous tubules by the efferent duct and testicular interstitium, respectively. Approximately 8 μL of a mixture containing rAAV1, HBSS and 4% Trypan blue solution was added to the testes of the recipient mice. Each injection filled 80–90% of the seminiferous tubules or testicular interstitium.

### Western immunoblotting analysis and silver staining analysis

Tissues were washed in 1 × PBS three times and then transferred to radioimmunoprecipitation assay (RIPA) solution (P0013C, Beyotime) containing 1 × protease inhibitor cocktail (11697498001, Roche) for cell/tissue lysis. The samples were spun at 12,000 ×g at 4°C for 30 min, and the supernatant was collected. Then, we added 5 × SDS loading buffer (P0015L, Beyotime) to the samples, thoroughly mixed them, and incubated them in boiling water for 10 min. For samples used to detect for protein expression of CLDN11, they should be incubated in water at 50°C for 10 minutes. The samples were concentrated via 5% sodium dodecyl sulfate‒polyacrylamide gel electrophoresis (SDS‒PAGE) at 80 V and separated via 10% SDS‒PAGE at 150 V. The isolated proteins were transferred to a polyvinylidene fluoride (PVDF) (#162e0177, Bio-Rad) membrane for 90 min at 250 mA, blocked with 5% skim milk in triethanolamine-buffered saline with Tween 20 (TBST) at room temperature for 2 h and incubated overnight with primary antibodies at 4°C. After being washed three times in TBST, the samples were incubated at room temperature with a secondary antibody for 2 h. Next, the PVDF membrane was washed in TBST three times, and the protein bands were exposed with ECL reagent (#180-501, Tannon). The primary antibodies used were anti-Cas9 (1:800, 14697T, Cell Signaling Technology); anti-Cre (1:1000, 15036S, Cell Signaling Technology); anti-β-ACTIN (1:1000, A1978, Sigma); anti-GAPDH (1:2000, 10494-1-AP, Proteintech); anti-SOX9 (1:1000, AB5535, Millipore); anti-CLDN11 (1:500, AF5364, Affinity); anti-DDX4 (1:2000, ab13840, Abcam); and anti-ZO-1 (1:1000, 33-9100, Thermo Fisher Scientific). The secondary antibodies used were HRP-conjugated anti-rabbit IgG (1:1000, ZB2301, Zhongshan Jinqiao Biotechnology) and HRP-conjugated anti-mouse IgG (1:1000, ZB2305, Zhongshan Jinqiao Biotechnology).

To detect the rAAV1 capsid protein, virus particles were lysed and concentrated via 5% sodium dodecyl sulfate‒polyacrylamide gel electrophoresis (SDS‒PAGE) at 80 V and separated via 10% SDS‒PAGE for gel electrophoresis at 150 V. We then performed silver staining of the gel as described in the instructions of the Fast Sliver Stain Kit (P0017S, Beyotime).

### Tissue section, cultured cell and whole-mount immunofluorescence

For whole-mount analysis, seminiferous tubules from adult (8-week-old) C57BL/6 mice were prepared as previously described with modifications^58^. Briefly, testis tubules were washed with 1 × PBS, fixed in 5 mL of 4% PFA for 4‒5 h at 4°C, and incubated sequentially with 5 mL of 25%, 50%, 75%, and 100% methanol at 4°C each for 5 min. Testis tubules were frozen in 100% methanol at -20°C for long-term storage. After rehydration of the seminiferous tubules through a decreasing ethanol gradient (100%, 75%, 50%, 25%), immunostaining was performed overnight at 4°C with the primary antibodies. After being washed with PBS three times, the seminiferous tubules were treated with secondary antibodies and stained with DAPI (1:500, D9542, Sigma‒Aldrich) at room temperature for 2 h. For tissue section immunofluorescence, the tissues were placed in 4% paraformaldehyde (PFA) (P6148, Sigma‒Aldrich) overnight and then embedded in optimal cutting temperature compound (4853, Sakura). Five-micron-thick sections were cut from pretreated tissues by a cryostat (Leica) and then blocked in phosphate-buffered saline (PBS) with 3% bovine serum albumin (BSA) at room temperature for 2 h. Then, the tissue sections were incubated at 4°C overnight with primary antibody diluted in 3% BSA. After being washed with PBS three times, the sections were treated with secondary antibodies and stained with DAPI (1:500, D9542, Sigma‒Aldrich). For cellular immunofluorescence, cultured SCs were plated on glass coverslips, and after culture or viral transfection, the cells were washed 3 times with PBS, fixed with 4% PFA for 30 min, permeabilized with 0.1% Triton X-100 (ST795, Beyotime) for 15 min and blocked in PBS supplemented with 3% BSA for 2 h, all at room temperature. Staining was performed as described above. The primary antibodies used were anti-SOX9 (1:500, AB5535, Millipore), anti-Androgen Receptor (1:200, ab133273, Abcam), anti-Ki67 (1:1000, ab15580, Abcam), anti-CLDN11 (1:400, AF5364, Affinity), anti-CLDN5 (1:300, A10207, ABclonal), anti-GATA-4 (1:500, sc-1237, Santa Cruz), anti-hPLZF (1:400, AF2944, R&D), anti-γ-H2Ax (1:500, ab11174, Abcam), and anti-PNA-rhodamine (1:1000, RL-1072, VECTOR LABORATORIES).

The secondary antibodies used were Cy™5 Donkey Anti-Rabbit IgG (H+L) (1:500, 711-175-152, Jackson ImmunoResearch); Rhodamine Red™-X (RRX) Donkey Anti-Rabbit IgG (H+L) (1:500, 711-295-152, Jackson ImmunoResearch); TRITC Donkey Anti-Goat IgG (H+L) (1:500, 705-025-147, Jackson ImmunoResearch); Alexa Fluor™488 Donkey Anti-Goat IgG (H+L) (1:500, A11055, Thermo Fisher Scientific); and Alexa Fluor™488 Goat Anti-Rabbit IgG (H+L) (1:500, A11034, Thermo Fisher Scientific). A confocal laser scanning microscope (LSM 800, Carl Zeiss, Germany) was used for observation and imaging.

### H&E staining

The testes were immobilized overnight in Hurtman’s fixative (H0290, Sigma‒Aldrich) for 24 h, dehydrated through an increasing ethanol gradient (70%, 80%, 90%, 95% and 100% ethanol, respectively), cleared in xylene, embedded in paraffin and sectioned (5 μm). The sections were deparaffinized via two xylene treatments (15 min each) and rehydrated through a decreasing ethanol gradient (100%, 90%, 80%, and 70% ethanol, respectively). After being washed in 1 × PBS three times, the sections were stained with hematoxylin (G1120, Solarbio) solution for 2.5 min, washed in running water for 10 min and then treated with 0.1% hydrochloric acid for 2 s. At this point, the sections were washed again and then stained with eosin (G1120, Solarbio) for 40 s. When the color of the sections became perceptible, the dyeing process was ended, and the sections were dehydrated through graded ethanol. The sections were subsequently treated twice with xylene (5 min each), sealed with neutral resin and finally photographed with a Nikon-112 microscope (Nikon, Japan).

### Transmission electron microscopy (TEM) analysis

The rAAV1-*iCre* and rAAV1-*Cldn11*-donor suspensions were mixed with 4% PFA at a 1:1 ratio to inactivate the virus and then sent to the Analysis and Testing Center of Nanjing Medical University for sample preparation. The virus sample was dropped onto a copper plate with a membrane and allowed to stand for 1 minute to remove excess liquid. The sample was then stained with 2% uranyl acetate to increase contrast for 1 minute. After drying at room temperature, it was observed using a transmission electron microscope (Tecnai G2 Spirit BioTwin, FEI, USA).

### PEM-seq assay

Six days after the transfection with rAAV1-*iCre* and rAAV1-*Cldn11*-donor, the *Cldn11^KO^-Rosa26^Cas9-tdTomato^* SCs (1 × 10^7^ in total from 8–10 mice) were collected and sent to Gene Medical Technology Co., Ltd. (Zhuhai, China) for primer-extension-mediated sequencing. The SCs from the *Cldn11^KO^-Rosa26^Cas9-tdTomato^* mice without treatment were used for the control library. Approximately 20 µg of genomic DNA was extracted and fragmented into 300–700 bp fragments by sonication. After removing the excess biotin-labeled primers, the biotin-labeled primers were used for primer extension amplification. The biotin-labeled single-stranded DNA was enriched with streptavidin magnetic beads, and the ends of the adapters containing random molecular barcodes were connected. Then, nested PCR was performed with a second set of primers to amplify the single-stranded DNA connected to the adapters. The products were subjected to tagged PCR and sequenced and analyzed on the MGI 2000 platform with 150 bp paired-end reads.

### Flow cytometric sorting

Testes from adult mice were kept in 1 × PBS, and the tunica albuginea was stripped under a stereoscope. The mass of the seminiferous tubules was stretched and incubated in DMEM (11965-084, Gibco) containing 1 mg/mL collagenase IV (C5138, Sigma‒Aldrich) for 5 min at 37°C with gentle shaking. The dispersed seminiferous tubules were washed with DMEM and then centrifuged at 300 ×g. Then, the seminiferous tubules were incubated in 0.25% trypsin-EDTA (25200-056, Gibco) containing 1 mg/mL deoxyribonuclease I (DN25, Sigma‒Aldrich) for 5 min at 37°C with gentle shaking, and the reaction was terminated with fetal bovine serum. The cell suspension was filtered with a 40 μm cell strainer and centrifuged at 600 ×g for 5 min at 4°C, after which the supernatant was discarded. Next, the cells were mixed well in DMEM and sorted by a BD FACSAria Fusion instrument. Following sorting, the cells were centrifuged at 600 ×g for 5 min at 4°C, and the genomic DNA was extracted from the sorted cells as described previously. To collect the sperm in the testes, the cells were stained with Hoechst (H3570, Life Technologies) for 20 min at 37°C and then sorted via flow cytometry. The sperm were collected via centrifugation, preserved in FERTIUP Cryoprotectant for mouse spermatozoa (KYD-001-EX, Cosmo Bio), and then stored in liquid nitrogen for subsequent intracytoplasmic single sperm microinjection (ICSI).

### Intracytoplasmic sperm injection (ICSI) and embryo transfer

6-week-old wild-type female Institute of Cancer Research (ICR) mice were intraperitoneally injected with PMSG (#190915, Ningbo Sansheng) to stimulate the development of antral follicles, and 46 h later, these mice were injected with hCG (#180428, Ningbo Sansheng). Sixteen hours after ovulatory hCG injection, the cumulus-oocyte complexes were surgically isolated from the fallopian tube, and digested with 0.5 mg/mL hyaluronidase (#H4272, Sigma‒Aldrich) for oocyte collection. The sperm heads were isolated and aspirated into the pipette. A piezopulse penetrated the membrane, and the sperm head was deposited into the ooplasm. The pipette was retracted while gently aspirating to seal the oocyte membrane. Then the oocytes were maintained in the potassium simplex optimization medium (KSOM, #MR-020P-5F, Millipore) medium at 37°C under 5% CO2 in air. Two-cell embryos were then transferred into the uterine horn of each pseudo-pregnant ICR females.

### Statistical analysis and image layout

Unless otherwise specified, at least three independent replicates were performed for each experiment. For the comparison of two sets of data, we employed two independent samples T test and Welch’s correction; when multiple sets of data are being compared, we adopted one-way ANOVA and Welch-Forsythe correction. In particular, we utilized repeated measures ANOVA and Geisser-Greenhouse correction to compare the data in Figure 1E. For the data sets where there was at least one group that does not follow a normal distribution, we used Kruskal-Wallis H test (Figure 2D, 2E, 5H,5I and S5E) or Mann-Whitney U test (Figure 1C, S5B and S5H) for data analysis. All the results obtained are expressed as the means ± SEMs. \**P* < 0.05, \*\**P* < 0.01, \*\*\**P* < 0.001, \*\*\*\**P* < 0.0001. *P* < 0.05 was considered to indicate a significant difference. The software used for data analysis was GraphPad Prism 8 (Version 8.0.2), and that for image layout was Adobe Illustrator 2022 (Version 26.0.1).

## Data availability

The data that support the findings of this study are available from the corresponding author upon reasonable request.

## Acknowledgments

We thank Dr. Xuejiang Guo for sharing the *Amh^Cre^* mice, Dr. Yiqiang Cui for assisting in constructing the plasmids used in this study, and all the members of the Wu Laboratory for their kind assistance. We thank *Springer Nature* editing service for help with manuscript grammar checking. The work was supported by the National Natural Science Foundation of China (32470896, 32270897 to X.W.).

## Author contributions

**Xin Wu** conceived the study. **Tao Zhang**, **Anhao Guo** and **Yuan Chen** bred the mice. **Tao Zhang**, **Hanben Wang**, **Anhao Guo**, **Yuan Chen** and **Lufan Li** performed most of the experiments. **Hanben Wang** and **Tao Zhang** designed the plasmids for virus synthesis and primers to verify the results of gene integration. **Tao Zhang** and **Xin Wu** prepared the figures. **Tao Zhang**, **Anhao Guo** and **Xin Wu** wrote the manuscript. The manuscript was reviewed by all authors.

## Graphical Abstract

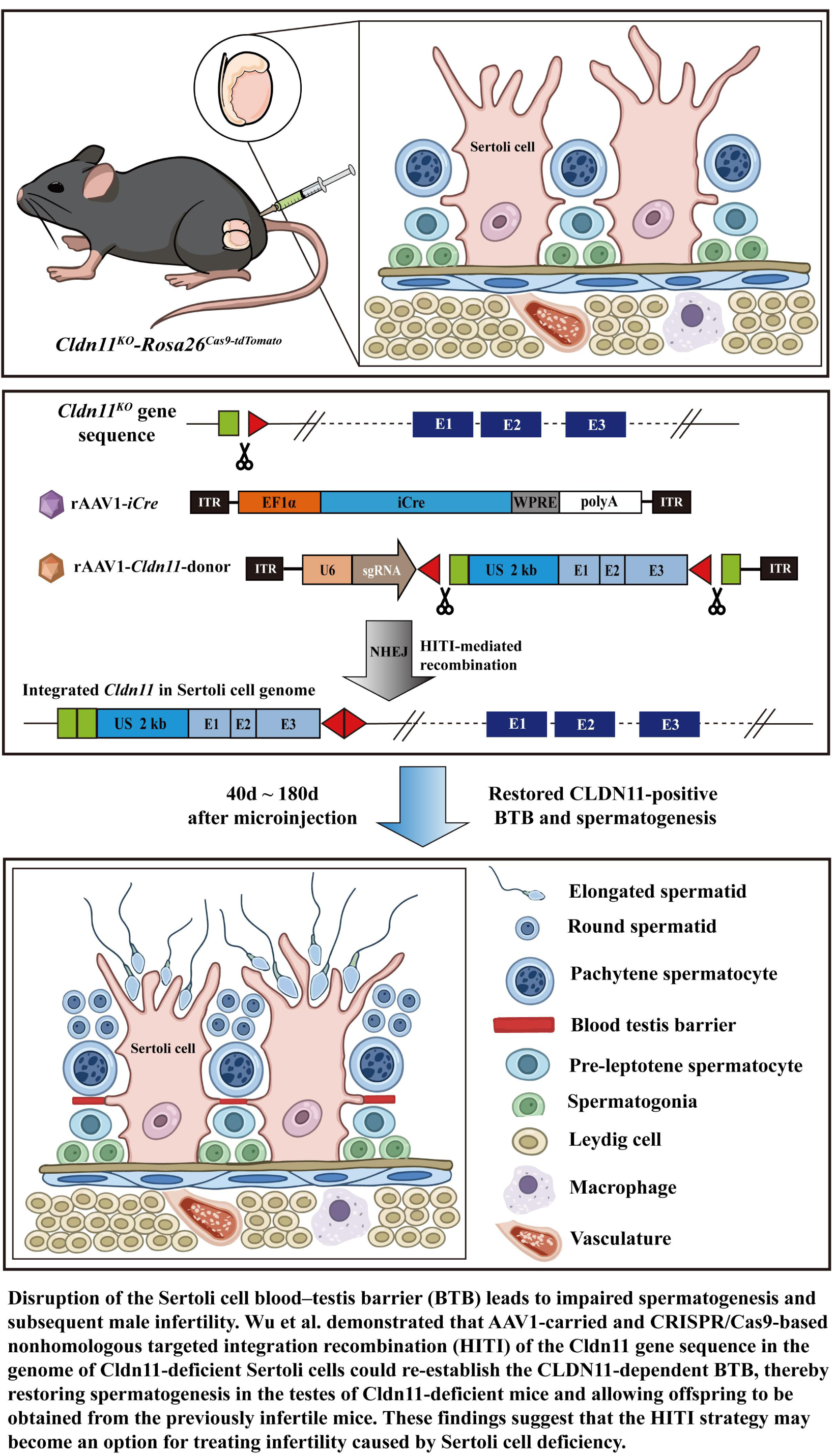

