## Supplemental materials for "Homology-independent targeted integration *in vivo* restores *Cldn11* deficiency in mouse Sertoli cells and spermatogenesis"

**Supplementary data**

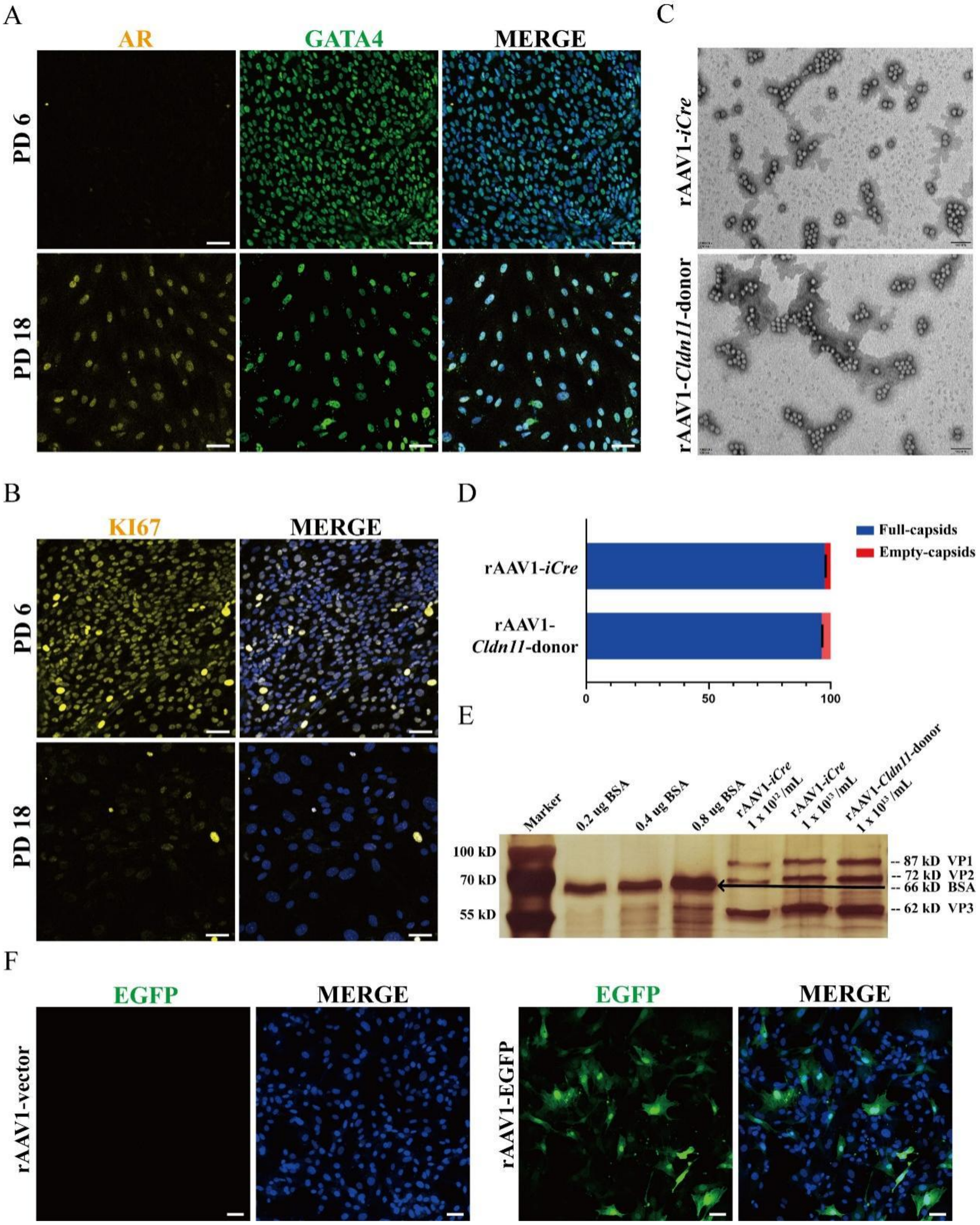


**Figure S1. Quality assurance of rAAV1-*iCre* & rAAV1-*Cldn11*-donor and *in vitro* culture of primary Sertoli cells and MEFs**

(A) Immunofluorescence of androgen receptor (AR, yellow) and GATA4 (green) in *Rosa26*$^{tomato\text{-}tdTomato}$ primary Sertoli cells cultured *in vitro*. Scale bars: 20 µm.

(B) Immunofluorescence staining of KI67 (yellow) in *Rosa26*$^{Cas9\text{-}tdTomato}$ primary Sertoli cells cultured *in vitro*. Scale bars: 20 µm.

(C) Representative transmission electron micrographs (TEMs) showing the plasmid packaging of rAAV1 particles. Scale bars: 100 nm.

(D) Ratios of full/empty capsids of rAAV1-*iCre* and rAAV1-*Cldn11*-donor.

(E) Silver staining of rAAV1-*iCre*, rAAV1-*Cldn11*-donor and bovine serum albumin (BSA) samples showing the ratios of the three capsid proteins of rAAV1.

(F) Immunofluorescence of EGFP (green) in MEFs transfected with rAAV1-EGFP *in vitro*. Scale bars: 20 μm.

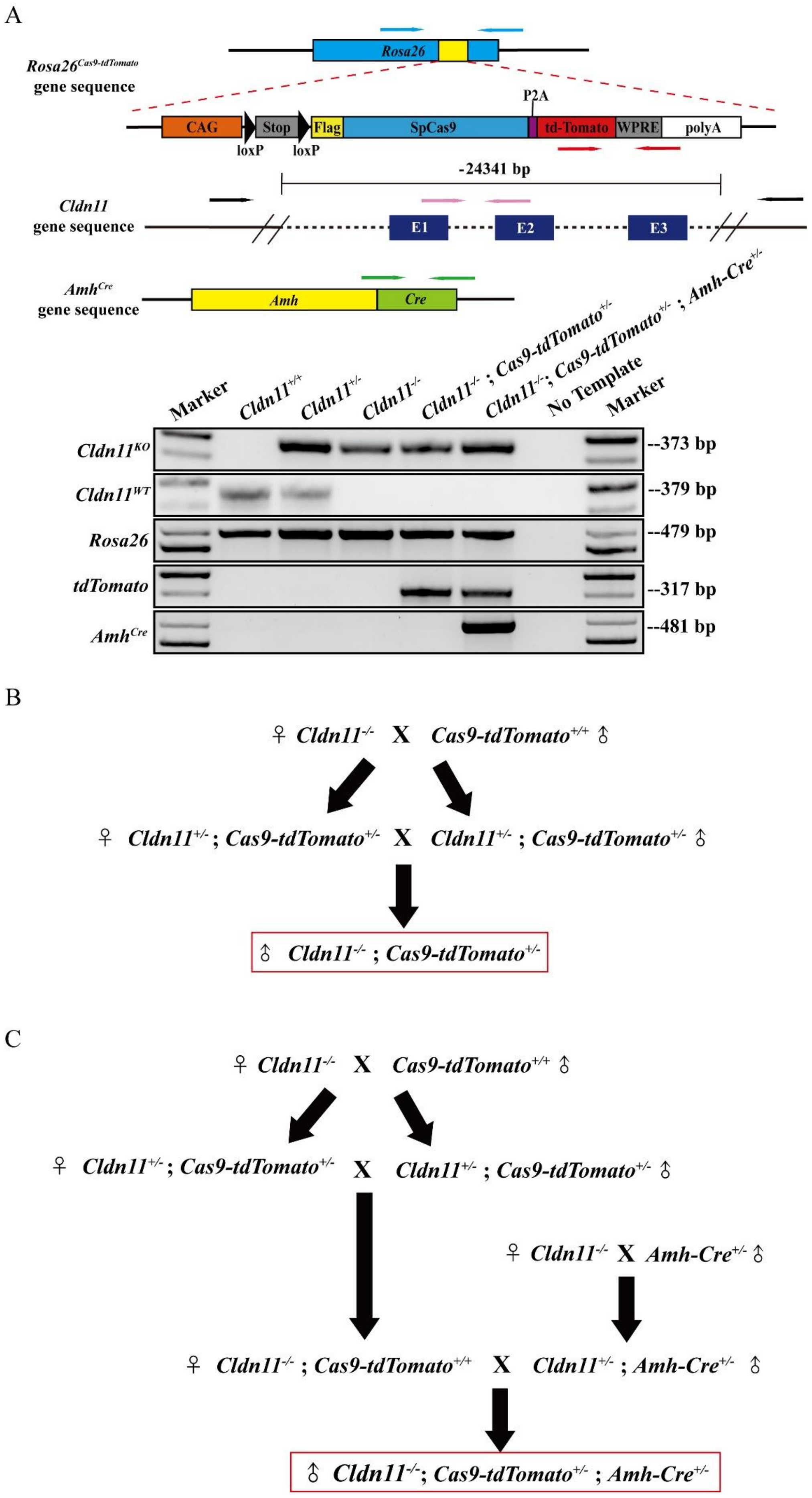
A
Rosa26
Rosa26Cas9-tdTomato gene sequence
CAG
Stop
Flag
SpCas9
P2A
td-Tomato
WPRE
polyA
loxP
loxP
-24341 bp
Cldn11 gene sequence
E1
E2
E3
AmhCre gene sequence
Amh
Cre
Marker
Cldn11+/+
Cldn11+/-
Cldn11-/-
Cldn11-/- ; Cas9-tdTomato+/-
Cldn11-/-; Cas9-tdTomato+/- ; Amh-Cre+/-
No Template
Marker
Cldn11KO
Cldn11WT
Rosa26
tdTomato
AmhCre
--373 bp
--379 bp
--479 bp
--317 bp
--481 bp
B
♀ Cldn11-/- X Cas9-tdTomato+/+ ♂
♀ Cldn11+/- ; Cas9-tdTomato+/- X Cldn11+/- ; Cas9-tdTomato+/- ♂
♂ Cldn11-/- ; Cas9-tdTomato+/-
C
♀ Cldn11-/- X Cas9-tdTomato+/+ ♂
♀ Cldn11+/- ; Cas9-tdTomato+/- X Cldn11+/- ; Cas9-tdTomato+/- ♂
♀ Cldn11-/- X Amh-Cre+/- ♂
♀ Cldn11-/- ; Cas9-tdTomato+/+ X Cldn11+/- ; Amh-Cre+/- ♂
♂ Cldn11-/-; Cas9-tdTomato+/- ; Amh-Cre+/-

**Figure S2. Genotyping and mating strategies of experimental mice**

(A) Schematic graphs of *Rosa26*$^{Cas9\text{-}tdTomato}$, *Cldn11*$^{KO}$ *and Amh*$^{Cre}$ transgenic mouse gene sequences and representative images of the genotyping results. The blue, red, black, pink and green arrows in the figure represent the *Rosa26, td-Tomato, Cldn11*$^{KO}$*, Cldn11*$^{WT}$ and *Amh*$^{Cre}$ primers, respectively.

(B) Schematic representation of the mating strategy used to obtain *Cldn11*$^{KO}$*-Rosa26*$^{Cas9\text{-}tdTomato}$ mice.

(C) Schematic representation of the mating strategy used to obtain *Cldn11*$^{KO}$*-Amh*$^{Cre}$*-Rosa26*$^{Cas9\text{-}tdTomato}$ mice.

A

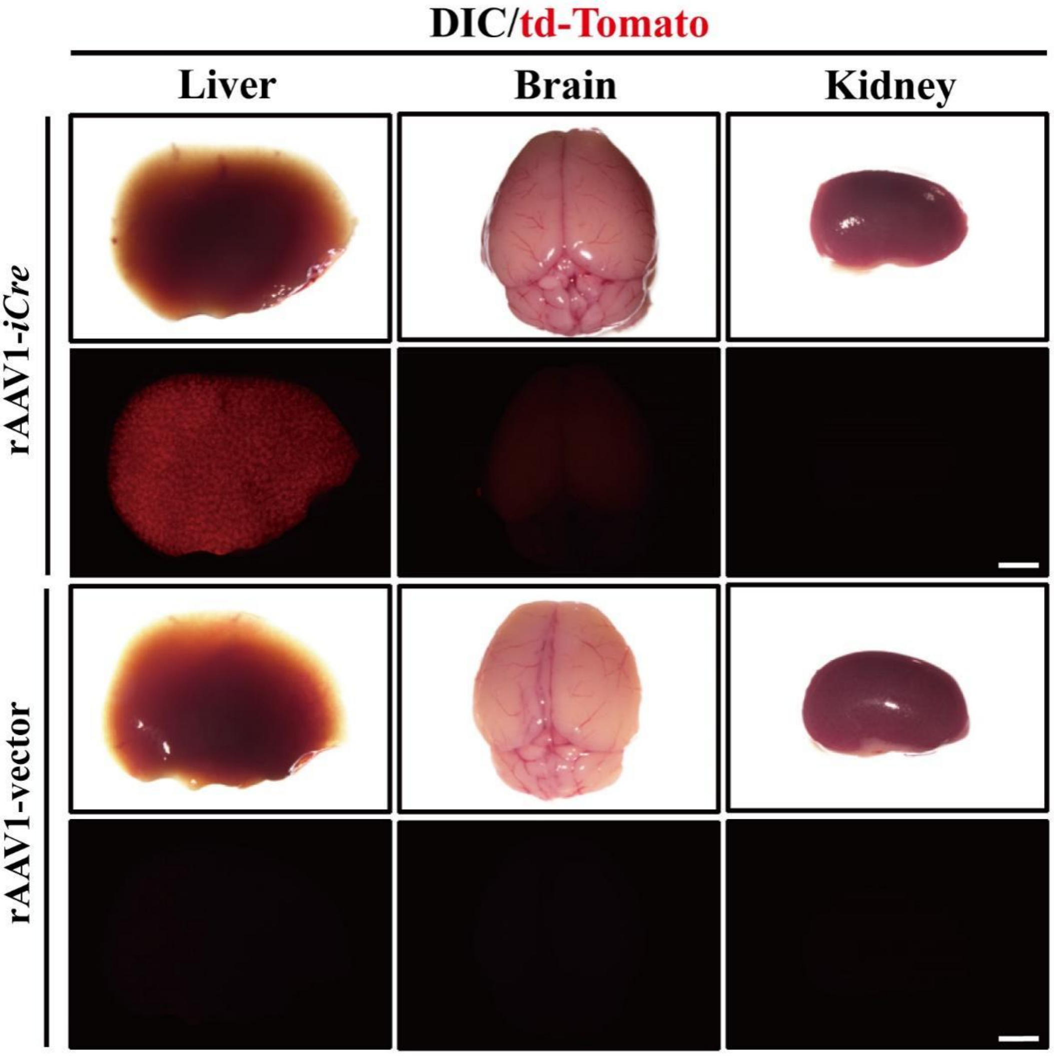


**Figure S3. Infection with rAAV1-*iCre* in various organs of *Rosa26*$^{Cas9\text{-}tdTomato}$ mice 40 days after transduction through tail vein injection**

(A) Representative liver, brain and kidney tissues from *Rosa26*$^{Cas9\text{-}tdTomato}$ mice under a fluorescence microscope 30 days after rAAV1-*iCre* transduction through the tail vein. The intensity of td-Tomato signal (red) in the organs is shown. Scale bars: 1 mm.

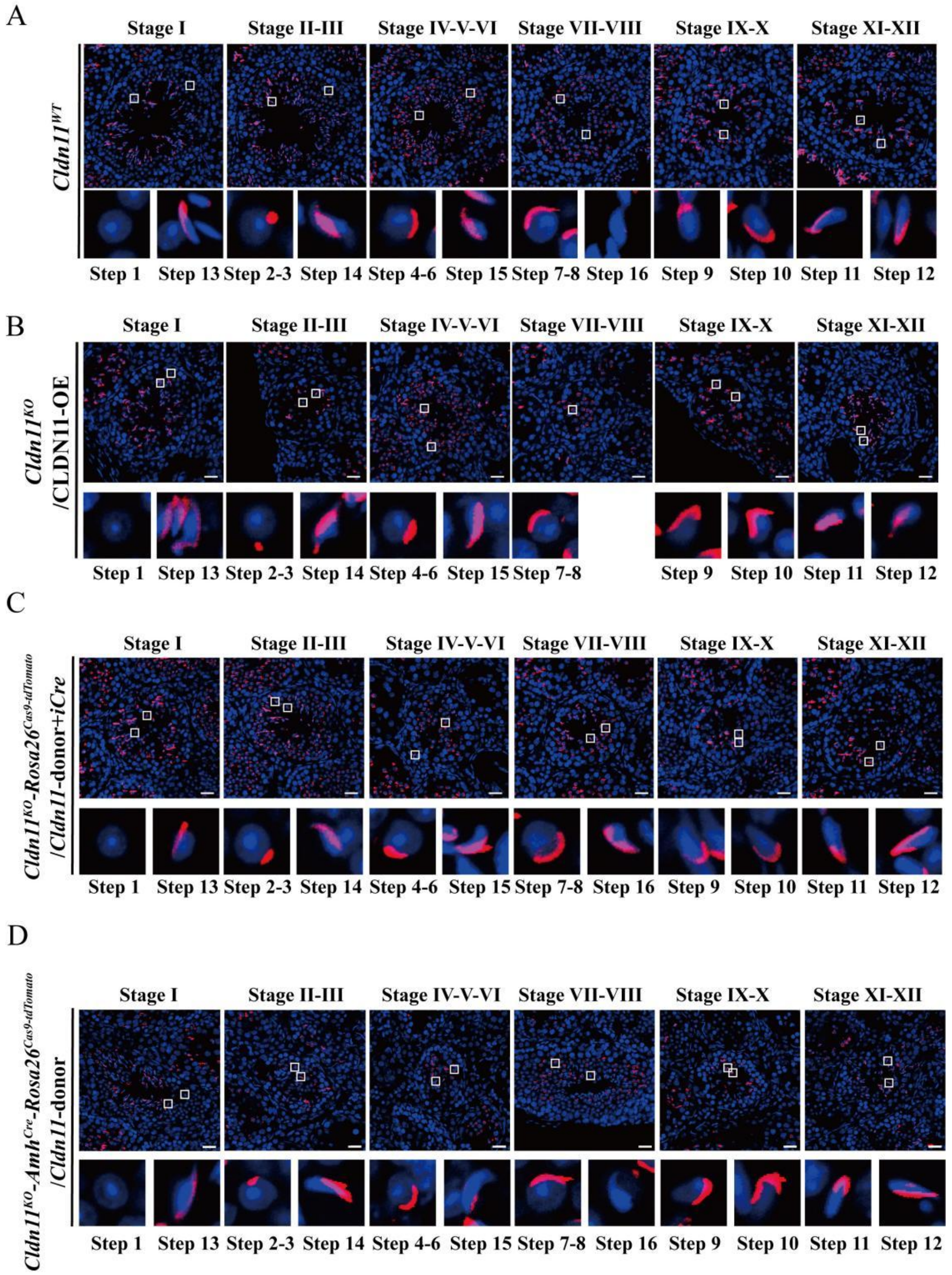


**Figure S4. Immunofluorescence of PNA in *Cldn11*$^{WT}$ testes and testes from the three treatment groups**

(A-D) PNA (red)- and DAPI (blue)-stained testis sections from *Cldn11*$^{WT}$ mice and the testes of the three treatment groups. The seminiferous tubule stage was identified on the basis of the acrosomal morphology marked by the PNA and the arrangement of spermatogenic cells, and the 16-step spermiogenesis process of round and elongated spermatid formation was identified by the PNA signal pattern. Scale bars: 20 μm.

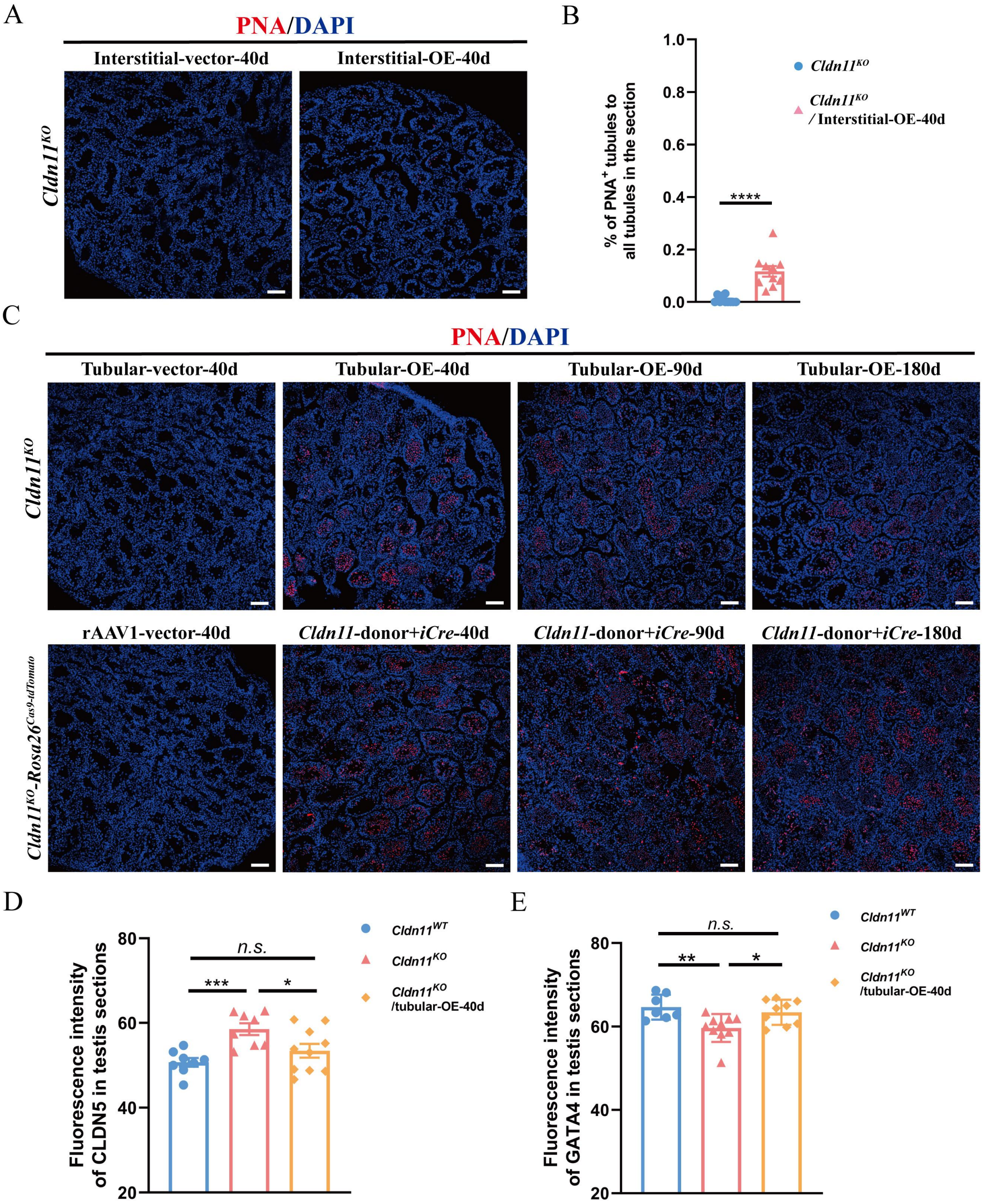
A
PNA/DAPI
Interstitial-vector-40d
Interstitial-OE-40d
Cldn11KO
B
1.0
0.8
0.6
0.4
0.2
0.0
% of PNA+ tubules to all tubules in the section
****
Cldn11KO
Cldn11KO / Interstitial-OE-40d
C
PNA/DAPI
Tubular-vector-40d
Tubular-OE-40d
Tubular-OE-90d
Tubular-OE-180d
Cldn11KO
rAAV1-vector-40d
Cldn11-donor+iCre-40d
Cldn11-donor+iCre-90d
Cldn11-donor+iCre-180d
Cldn11KO-Rosa26Cas9-tdTomato
D
80
60
40
20
Fluorescence intensity of CLDN5 in testis sections
n.s.
***
*
Cldn11WT
Cldn11KO
Cldn11KO /tubular-OE-40d
E
80
60
40
20
Fluorescence intensity of GATA4 in testis sections
n.s.
**
*
Cldn11WT
Cldn11KO
Cldn11KO /tubular-OE-40d

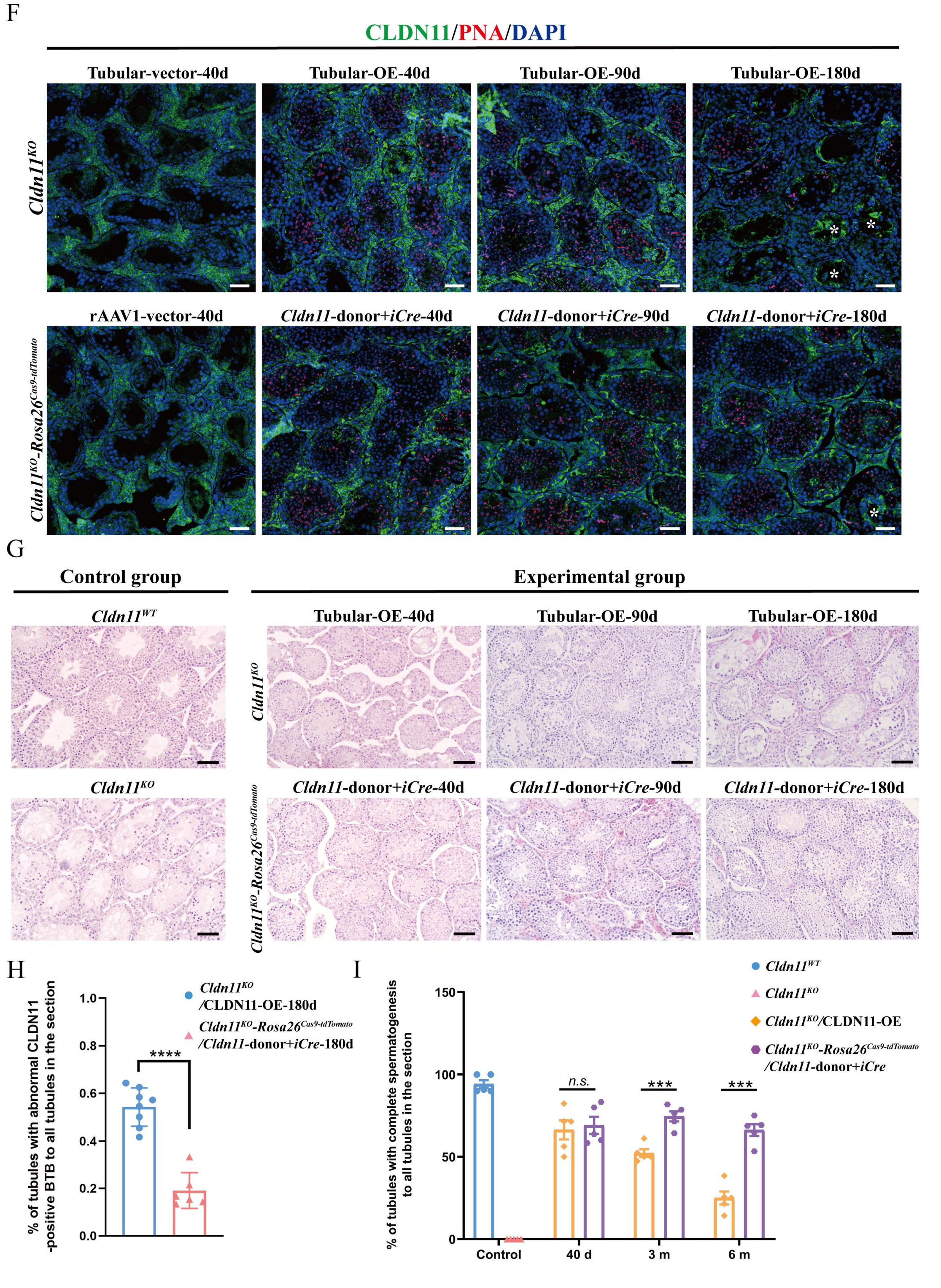


**Figure S5. Comparison of the duration of spermatogenesis rescue among the different treatment groups**

(A) Immunofluorescence of PNA (red) in *Cldn11*$^{KO}$ testes with rAAV1-*Cldn11*-donor transduced through the testicular interstitium for 40 days. rAAV1-vector transduction was used as a control. Scale bars: 20 μm.

(B) The percentage of $PNA^+$ tubules in *Cldn11*$^{KO}$ testes transduced with the rAAV1-*Cldn11*-donor through the testicular interstitium for 40 days compared with that in *Cldn11*$^{KO}$ testes transduced with the rAAV1-vector. Mann-Whitney U test was used for statistical analyses and $Z$= -3.862.

(C) Immunofluorescence of PNA (red) in *Cldn11*$^{KO}$ testes and *Cldn11*$^{KO}$*-Rosa26*$^{Cas9\text{-}tdTomato}$ testes with rAAV1 transduced through seminiferous tubules for different lengths of time. rAAV1-vector transduction was used as a control. Scale bars: 20 μm.

(D) Histograms of the fluorescence intensity of CLDN5 in *Cldn11*$^{WT}$ and *Cldn11*$^{KO}$ testes and *Cldn11*$^{KO}$ testes transduced with the rAAV1-*Cldn11*-donor for 40 days. One-way ANOVA was used for statistical analyses and $F$=7.222.

(E) Histograms of the fluorescence intensity of GATA4 in *Cldn11*$^{WT}$ and *Cldn11*$^{KO}$ testes and *Cldn11*$^{KO}$ testes transduced with the rAAV1-*Cldn11*-donor for 40 days. Kruskal-Wallis H test was used for statistical analyses and $H$=8.971.

(F) Immunofluorescence of CLDN11 (green) and PNA (red) staining in *Cldn11*$^{KO}$ testes and *Cldn11*$^{KO}$*-Rosa26*$^{Cas9\text{-}tdTomato}$ testes with rAAV1 transduced through seminiferous tubules for different lengths of time. rAAV1-vector transduction was used as a control. Asterisks indicate tubules with abnormal CLDN11-positive BTB. Scale bars: 20 μm.

(G) Testis histology revealed spermatogenesis in *Cldn11*$^{KO}$ testes and *Cldn11*$^{KO}$*-Rosa26*$^{Cas9\text{-}tdTomato}$ testes with rAAV1 transduced through seminiferous tubules for different lengths of time, compared with *Cldn11*$^{WT}$ and *Cldn11*$^{KO}$ testes. Scale bars: 50 μm.

(H) Percentage of tubules with abnormal CLDN11-positive BTB 180 days after transduction with rAAV1-*Cldn11*-donor in *Cldn11*$^{KO}$ testes and transduction with rAAV1-*Cldn11*-donor and rAAV1-*iCre* in *Cldn11*$^{KO}$*-Rosa26*$^{Cas9\text{-}tdTomato}$ testes through the seminiferous tubules. Mann-Whitney U test was used for statistical analyses and $Z$= -3.098.

(I) Percentage of tubules with complete spermatogenesis in *Cldn11*$^{KO}$ testes transduced with rAAV1-*Cldn11*-donor and *Cldn11*$^{KO}$*-Rosa26*$^{Cas9\text{-}tdTomato}$ testes transduced with rAAV1-*Cldn11*-donor and rAAV1-*iCre* through seminiferous tubules for different lengths of time. Two independent samples T test was used for statistical analyses.

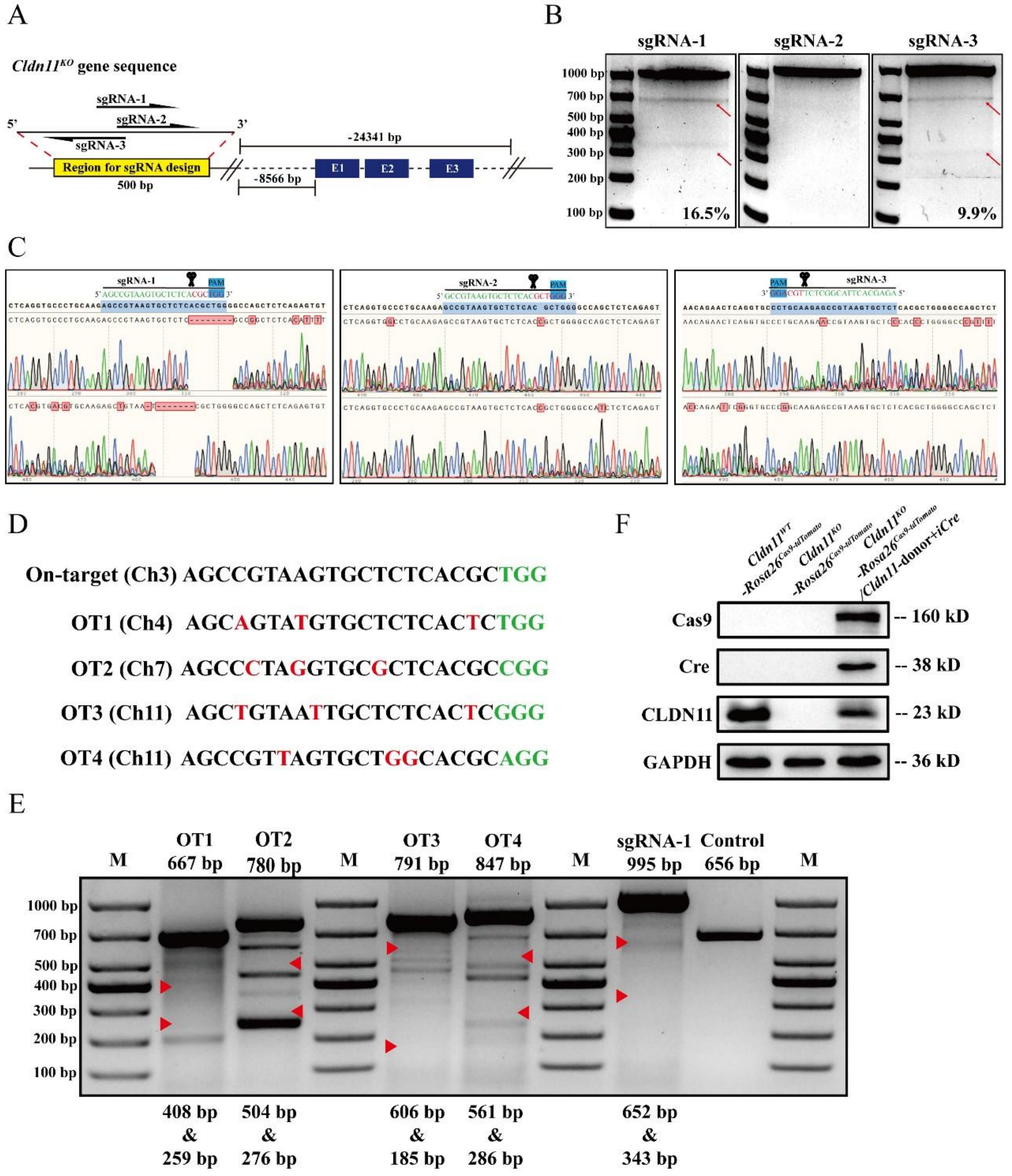


**Figure S6. Detection of the cleavage activity of three sgRNAs and off-target detection of sgRNA-1**

(A) Schematic of the design of the three sgRNAs.

(B) Assessment of editing efficiency at target sites in primary cultured SCs via T7EI assays.

(C) Graph of Sanger sequencing showing the Cas9 cleavage of DNA guided by three sgRNAs.

(D) Potential off-target sites predicted by the ATUM gRNA design tool. Nucleotide mismatches are indicated in red; the PAM is indicated in green.

(E) Assessment of editing efficiency at possible off-target sites in primary cultured Sertoli cells via T7EI assays. The red arrows indicate the locations of the predicted bands.

(F) Western blotting verified Cas9, Cre and CLDN11 protein expression in *Cldn11*$^{KO}$-*Rosa26*$^{Cas9\text{-}tdTomato}$ SCs with HITI strategy-mediated *Cldn11* integration compared with that in *Cldn11*$^{WT}$-*Rosa26*$^{Cas9\text{-}tdTomato}$ and *Cldn11*$^{KO}$-*Rosa26*$^{Cas9\text{-}tdTomato}$ SCs. GAPDH was used as the protein loading control.

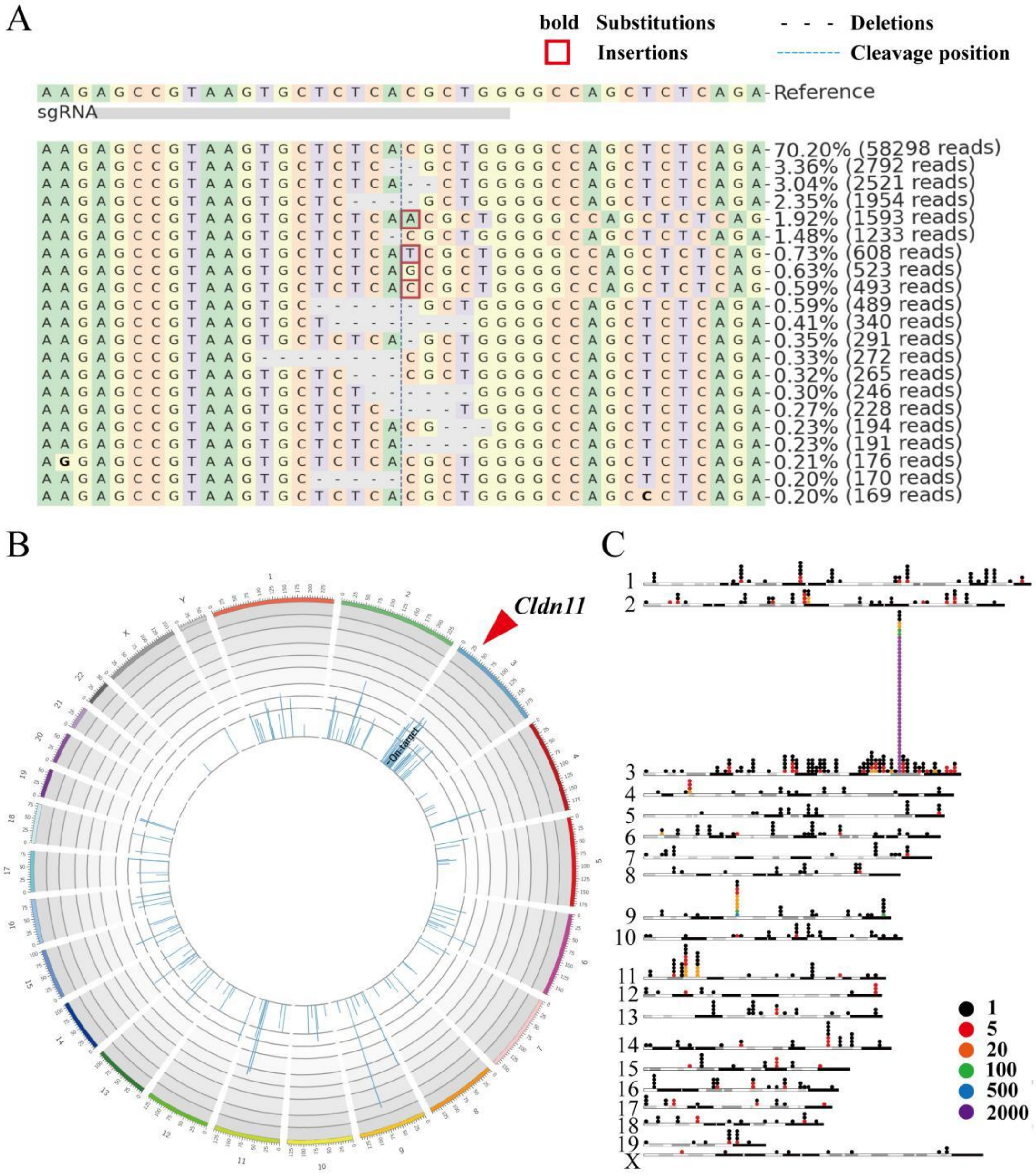


**Figure S7. Primer-extension-mediated sequencing (PEM-seq) on on-target editing and off-target hotspots of sgRNA-1/Cas9 mediated HITI in SCs**

(A) Visualization of editing in the on-target site. The reference sequence (upper panel) contains the target editing site, and the reads (lower panel) are compared to the upper reference sequence. Nucleotides are indicated by different colors respectively (A: green; C: pink; G: yellow; T: purple). The vertical blue dotted line indicates the DSB position, bold letters indicate replaced sequences, red rectangles indicates inserted sequences, and horizontal dotted lines indicates deleted sequences. The graph shows the cases with a percentage ≥ 0.2% and does not include reads of translocations.

(B) Circos plots of sgRNA-1/Cas9 libraries. Chromosomes are shown with centromere to telomere in the clockwise direction, and the numbers on the outer circle represent the numbers of chromosome and the scale represents the coordinate of the chromosome (5 Mbp per scale). Red arrow indicates the sgRNA-1/Cas9 cleavage site (On-target). The blue bar in the inner circle counts the numbers of the translocation events (in logarithms) on the chromosome interval. The curve in the inner circle connects from the on-target locus to the off-target locus, while no curves shown indicates that no off-target translocations are detected.

(C) Visualization of global captured editing events. The horizontal bars in the figure represent chromosomes binned into 2 Mb regions, and the colored circles represent the numbers of editing events.

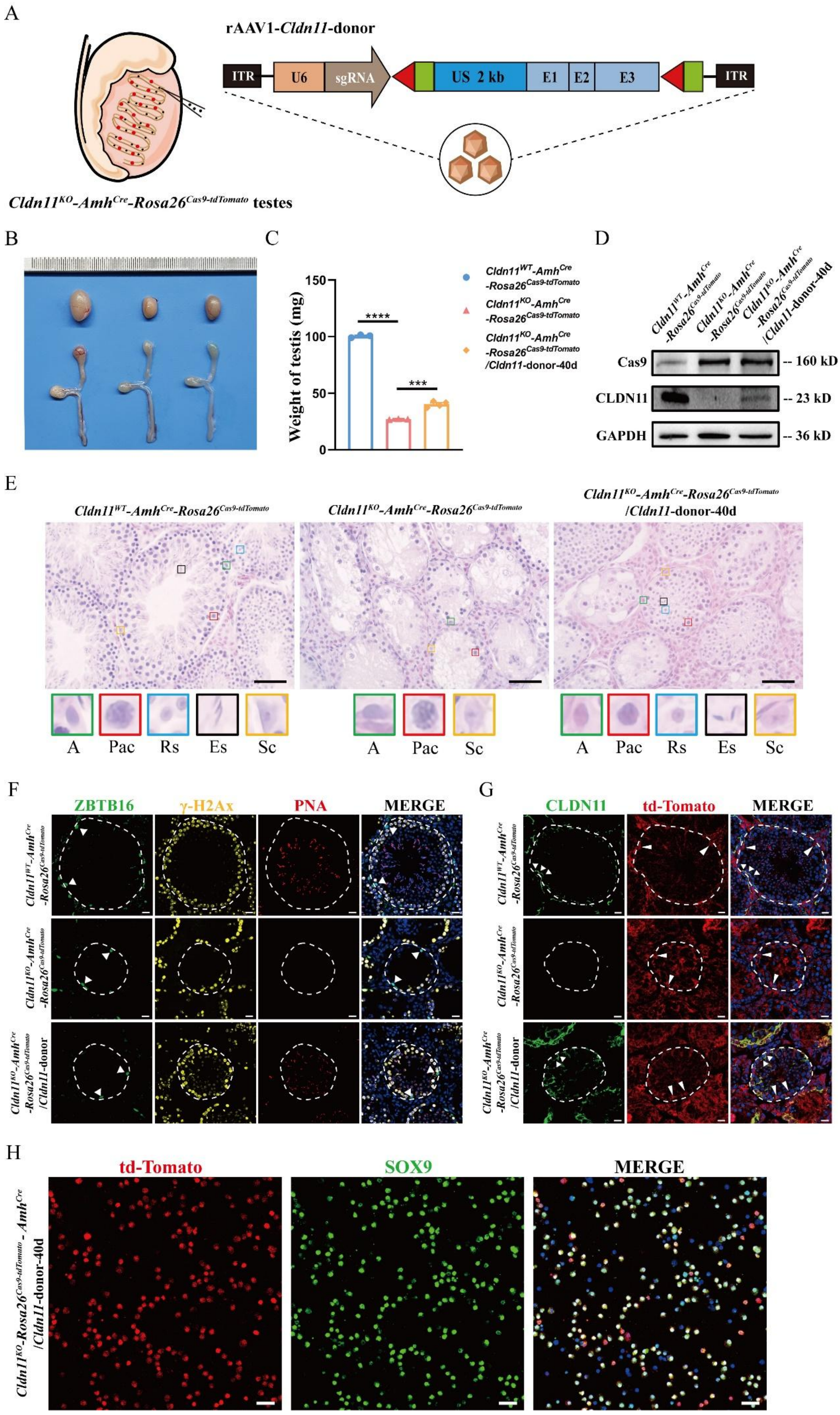
A
rAAV1-Cldn11-donor
ITR
U6
sgRNA
US 2 kb
E1
E2
E3
ITR
Cldn11KO-AmhCre-Rosa26Cas9-tdTomato testes
B
C
Weight of testis (mg)
150
100
50
0
****
***
Cldn11WT-AmhCre-Rosa26Cas9-tdTomato
Cldn11KO-AmhCre-Rosa26Cas9-tdTomato
Cldn11KO-AmhCre-Rosa26Cas9-tdTomato/Cldn11-donor-40d
D
Cas9
-- 160 kD
CLDN11
-- 23 kD
GAPDH
-- 36 kD
E
A
Pac
Rs
Es
Sc
F
ZBTB16
γ-H2Ax
PNA
MERGE
Cldn11KO-AmhCre-Rosa26Cas9-tdTomato/Cldn11-donor
G
CLDN11
td-Tomato
MERGE
H
td-Tomato
SOX9
MERGE
Cldn11KO-Rosa26Cas9-tdTomato-AmhCre/Cldn11-donor-40d

**Figure S8. *Cldn11* genomic integration via HITI specially in SCs rescues spermatogenesis in *Cldn11*-deficient mouse testes**

(A) Schematic graphs of testicular tubular injection of rAAV1-*Cldn11*-donor for validation of HITI of the *Cldn11* donor sequence specifically in Sertoli cells.

(B) Representative morphological image of testes and epididymides from *Cldn11*$^{WT}$-*Amh*$^{Cre}$-*Rosa26*$^{Cas9\text{-}tdTomato}$ and *Cldn11*$^{KO}$-*Amh*$^{Cre}$-*Rosa26*$^{Cas9\text{-}tdTomato}$ mice and *Cldn11*$^{KO}$-*Amh*$^{Cre}$-*Rosa26*$^{Cas9\text{-}tdTomato}$ mice 40 days after tubular injection with rAAV1-*Cldn11*-donor.

(C) Comparison of the testis weights of *Cldn11*$^{WT}$-*Amh*$^{Cre}$-*Rosa26*$^{Cas9\text{-}tdTomato}$ mice, *Cldn11*$^{KO}$-*Amh*$^{Cre}$-*Rosa26*$^{Cas9\text{-}tdTomato}$ mice *and* *Cldn11*$^{KO}$-*Amh*$^{Cre}$-*Rosa26*$^{Cas9\text{-}tdTomato}$ mice with tubular injection of rAAV1-*Cldn11*-donor ($n \geq 3$). One-way ANOVA was used for statistical analyses and $F$=1442.594.

(D) Western blotting verified Cas9 and CLDN11 protein expression in *Cldn11*$^{KO}$-*Amh*$^{Cre}$-*Rosa26*$^{Cas9\text{-}tdTomato}$ mouse testes subjected to tubular injection of rAAV1-*Cldn11*-donor compared with *Cldn11*$^{WT}$-*Amh*$^{Cre}$-*Rosa26*$^{Cas9\text{-}tdTomato}$ and *Cldn11*$^{KO}$-*Amh*$^{Cre}$-*Rosa26*$^{Cas9\text{-}tdTomato}$ mouse testes. GAPDH was used as the loading control.

(E) Extensive spermatogenesis in *Cldn11*$^{KO}$-*Amh*$^{Cre}$-*Rosa26*$^{Cas9\text{-}tdTomato}$ mouse testes with HITI strategy-mediated *Cldn11* integration compared with *Cldn11*$^{WT}$-*Amh*$^{Cre}$-*Rosa26*$^{Cas9\text{-}tdTomato}$ and *Cldn11*$^{KO}$-*Amh*$^{Cre}$-*Rosa26*$^{Cas9\text{-}tdTomato}$ testes. The lower panels with colored frames show the types of cells found in the mouse testes. A, type A spermatogonia; Pac, pachytene spermatocyte; Rs, round spermatid; Es, elongated spermatid; Sc, Sertoli cell. Scale bars: 50 μm.

(F) Immunofluorescence of ZBTB16 (green), γ-H2Ax (yellow) and PNA (red) in *Cldn11*$^{WT}$-*Rosa26*$^{Cas9\text{-}tdTomato}$ testes, *Cldn11*$^{KO}$-*Rosa26*$^{Cas9\text{-}tdTomato}$ testes and *Cldn11*$^{KO}$-*Rosa26*$^{Cas9\text{-}tdTomato}$ testes with HITI strategy-mediated *Cldn11* integration. Scale bars: 20 μm.

(G) Immunofluorescence of CLDN11 (green) in *Cldn11*$^{WT}$-*Amh*$^{Cre}$-*Rosa26*$^{Cas9\text{-}tdTomato}$ testes, *Cldn11*$^{KO}$-*Amh*$^{Cre}$-*Rosa26*$^{Cas9\text{-}tdTomato}$ testes and *Cldn11*$^{KO}$-*Amh*$^{Cre}$-*Rosa26*$^{Cas9\text{-}tdTomato}$ testes with HITI strategy-mediated *Cldn11* integration, shown by small arrows. Scale bars: 20 μm.

(H) Immunofluorescence of SOX9 (green) and td-Tomato (red) in SCs collected via flow cytometry from *Cldn11*$^{KO}$-*Amh*$^{Cre}$-*Rosa26*$^{Cas9\text{-}tdTomato}$ testes with HITI strategy-mediated *Cldn11* integration. Scale bars: 20 μm.

**Figure 1. F**

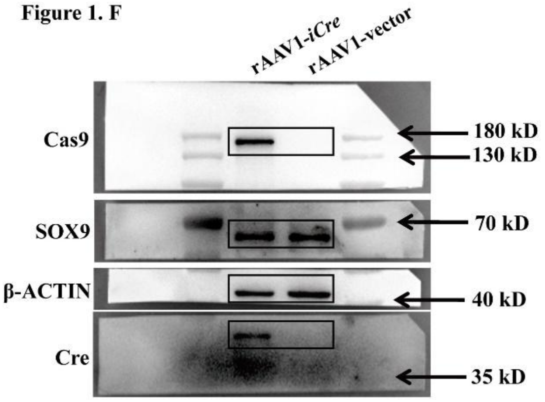


**Figure 3. A**

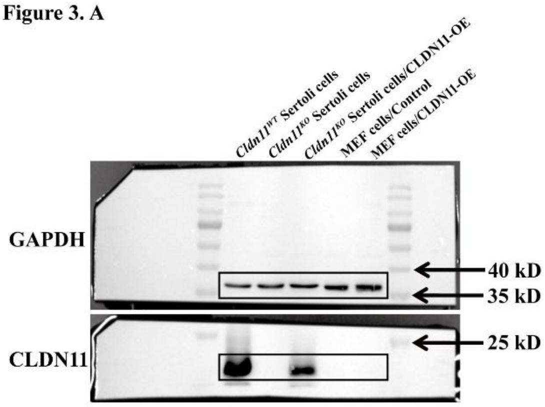


**Figure 3. E**

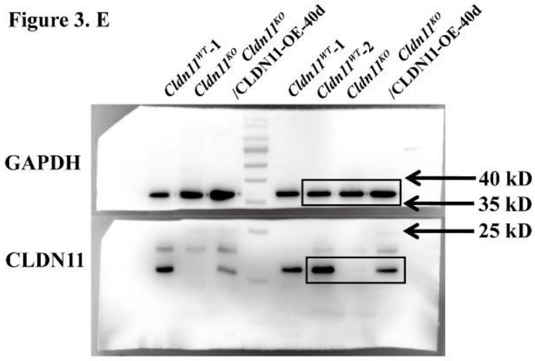


**Figure 5. D**

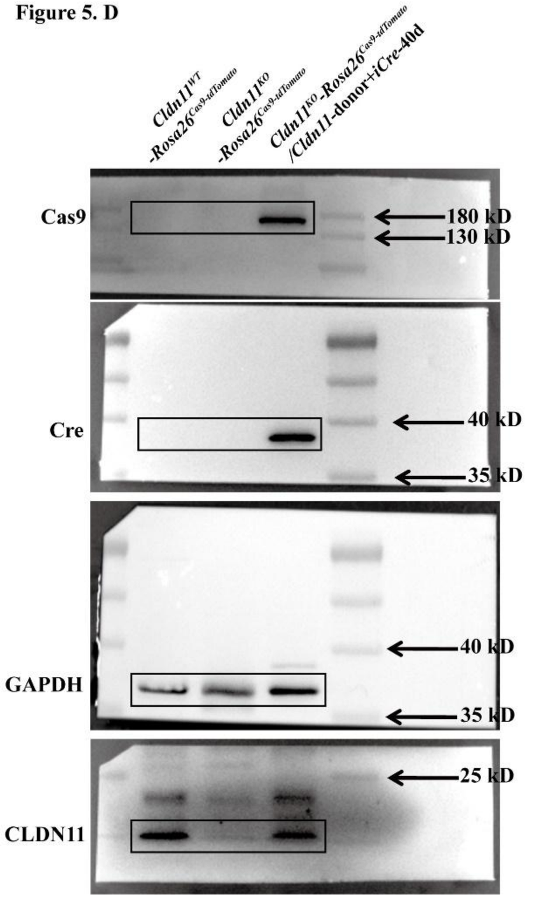


**Figure 6. C**

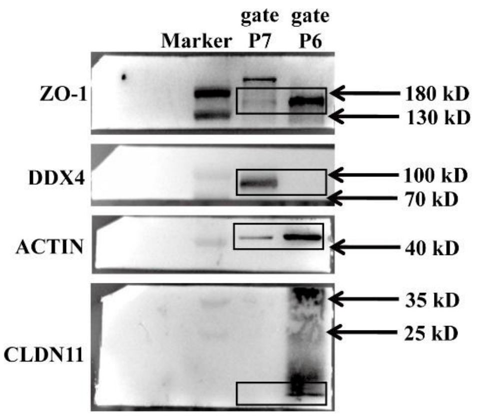

Figure S6. F

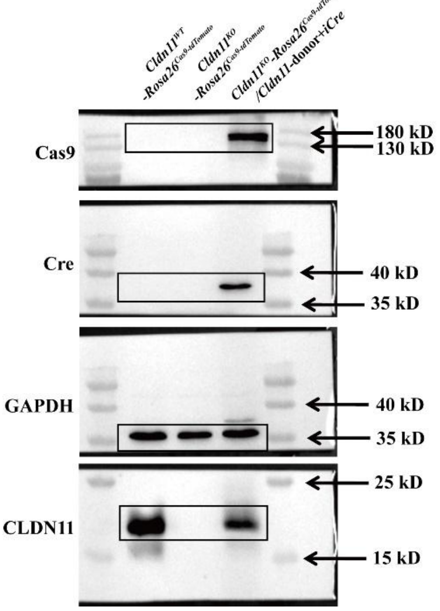


Figure S8. D

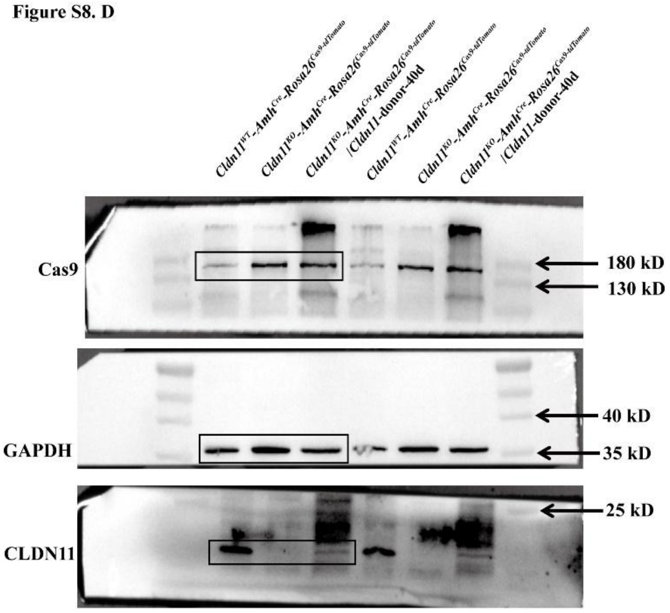


**Figure S9. Unprocessed images of western blotting**

PVDF membranes were cut into several small pieces to incubate with different antibodies for immunoblotting. Black boxes indicate images showed in relevant figures.

**Table S1: Key resources used in the research**

**Primers for genotyping**

| Gene | | Sequence | Size of product sequence |
|---|---|---|---|
| *Cldn11* | Forward-1<br>Reverse-1 | 5'-ATGCTGGGAACAGAACTCAGGTG-3'<br>5'-TCATCCCAAAACCAGGGACTG-3' | WT: 24714 bp<br>KO: 373 bp |
| | Forward-2<br>Reverse-2 | 5'-TACCACACTAGGTTGAGTGTGGC-3'<br>5'-CAACAAGGTCACCTGTGATGCTG-3' | WT: 379 bp<br>KO: 0 bp |
| *Rosa26*$^{Cas9\text{-}tdTomato}$ | Rosa26-Forward<br>Rosa26-Reverse | 5'-CCCAAAGTCGCTCTGAGTTGTTA-3'<br>5'-TCGGGTGAGCATGTCTTTAATCT-3' | 479 bp |
| | tdTomato-Forward<br>WPRE-Reverse | 5'-CGGCATGGACGAGCTGTACAAG-3'<br>5'-TCAGCAAACACAGTGCACACCAC-3' | 317 bp |
| *Amh-Cre* | Transgene-Forward<br>Transgene-Reverse | 5'-GCCTGCATTACCGGTCGATGC-3'<br>5'-CAGGGTGTTATAAGCAATCCC-3' | 481 bp |
| *Sry* | Forward<br>Reverse | 5'-AGATCTTGATTTTTAGTGTTC-3'<br>5'-TGCAGCTCTACTCCAGTCTTG-3' | 460 bp |

**Primers for verification of 5'- and 3'- integration generated by HITI strategy**

| Name | | Sequence | Size of product sequence |
|---|---|---|---|
| primers for 5'-integration | Forward<br>Reverse | 5'-GTATAGAGAGTCTGAGAAAGGAGGATTG-3'<br>5'-TGAGAATGGCAAACACTTTATTACTGTTA-3' | 446 bp |
| primers for 3'-integration | Forward<br>Reverse | 5'-CTGTCTCCATTCTGTTAGAGACCATG-3'<br>5'-TGTTCCCCCAATAGCTTCCTCC-3' | 576 bp |

**Sequences of target site and potential off-target site**

| Name | Sequence | Note |
|---|---|---|
| off-target 1 (OT1) | 5'-AGCAGTATGTGCTCTCACTCTGG-3' | Letters labeled in red represent incorrect bases in off-target site sequences (up to 3-base mismatches) |
| off-target 2 (OT2) | 5'-AGCCCTAGGTGCGCTCACGCCGG-3' | |
| off-target 3 (OT3) | 5'-AGCTGTAATTGCTCTCACTCGGG-3' | |
| off-target 4 (OT4) | 5'-AGCCGTTAGTGCTGGCACGCAGG-3' | |
| target site | 5'-AGCCGTAAGTGCTCTCACGCTGG-3' | |

**Primers for T7 Endonuclease I cleavage assay of the on-target and potential off-target sites**

| Name | | Sequence | Size of product sequence |
|---|---|---|---|
| primers for T7-OT1 | Forward<br>Reverse | 5'-GCCTTTGTTTAGGCTGGGTTCT-3'<br>5'-CAATGAGTCCCTTCTACTCCTGACT-3' | 667 bp |
| primers for T7-OT2 | Forward<br>Reverse | 5'-AGAAGTCGATCAGAGTGCGCT-3'<br>5'-AGTAGAAAGGGCTTTGTCCGC-3' | 780 bp |
| primers for T7-OT3 | Forward<br>Reverse | 5'-TGGCTTCCCTCTGTGATAGACT-3'<br>5'-GGGAGGGGAGAGAAGAGATAGCT-3' | 791 bp |
| primers for T7-OT4 | Forward<br>Reverse | 5'-CTCCTGACTTTCCTCCCCTTTAC-3'<br>5'-ACTACTTGGACAACAGAGGCAAG-3' | 847 bp |
| primers for T7-target site | Forward<br>Reverse | 5'-GTATAGAGAGTCTGAGAAAGGAGGATTG-3'<br>5'-TAGACCTGAGTATCAACTGGCACAA-3' | 995 bp |

**Primers for Integration Assay in offsprings**

| Name | | Sequence | Size of product sequence |
|---|---|---|---|
| U6 | Forward<br>Reverse | 5'-GCCTATTTCCCATGATTCCTTC-3'<br>5'-CCTTTCCACAAGATATATAAAGCCA-3' | 235 bp |
| 5'-ITR | Forward<br>Reverse | 5'-AGTGGCCAACTCCATCACTA-3'<br>5'-GTTACGGTAAGCATATGATAGTCC-3' | 240 bp |
| 3'-ITR | Forward<br>Reverse | 5'-TGTCTCCAAACCAGCGTGA-3'<br>5'-GGTGTGAAATACCGCACAGAT-3' | 242 bp |
| Region of exons | Forward<br>Reverse | 5'-CCGAAAAATGGACGAACTGG-3'<br>5'-GTACAGCGAGTAGCCAAAGC-3' | 355 bp |

**Antibodies used in this research**

| REAGENT or RESOURCE | SOURCE and IDENTIFIER | Dilution for WB | Dilution for IF |
|---|---|---|---|
| **Primary antibodies** | | | |
| Mouse Monoclonal anti-Cas9(*S. pyogenes*) | Cat#14697T, Cell Signaling Technology | 1:800 | |
| Rabbit Monoclonal anti-Cre Recombinase | Cat#15036S, Cell Signaling Technology | 1:1000 | |
| Mouse Monoclonal anti-β-ACTIN | Cat#A1978, Sigma | 1:5000 | |
| Rabbit Polyclonal anti-GAPDH | Cat#10494-1-AP, Proteintech | 1:2000 | |
| Rabbit Polyclonal anti-SOX9 | Cat#AB5535, Millipore | 1:1000 | 1:500 |
| Rabbit Polyclonal anti-CLDN11 | Cat#AF5364, Affinity | 1:500 | 1:400 |
| Rabbit Polyclonal anti-DDX4 | Cat#ab13840, Abcam | 1:2000 | |
| Mouse Monoclonal anti-ZO-1 | Cat#33-9100, Thermo Fisher Scientific | 1:1000 | |
| Rabbit Polyclonal anti-CLDN5 | Cat#A10207, ABclonal | | 1:300 |
| Goat Polyclonal anti-GATA-4 | Cat#sc-1237, Santa Cruz | | 1:500 |
| Goat Polyclonal anti-hPLZF (ZBTB16) | Cat#AF2944, R&D | | 1:400 |
| Rabbit Polyclonal anti-γ-H2Ax | Cat#ab11174, Abcam | | 1:500 |
| Rabbit polyclonal anti-Ki67 | Cat#ab15580, Abcam | | 1:1000 |
| Rabbit Monoclonal Anti-Androgen Receptor | Cat#ab133273, Abcam | | 1:200 |
| PNA, Rhodamine | Cat#RL-1072, VECTOR LABORATORIES | | 1:1000 |
| **Secondary antibodies** | | | |
| HRP-conjugated anti-rabbit IgG | Cat#ZB2301, Zhongshan Jinqiao biotechnology | 1:1000 | |
| HRP-conjugated anti-mouse IgG | Cat#ZB2305, Zhongshan Jinqiao biotechnology | 1:1000 | |
| Rhodamine Red™-X (RRX) AffiniPure Donkey Anti-Rabbit IgG (H+L) | Cat#711-295-152, Jacksonimmun | | 1:500 |
| Cy™5 AffiniPure Donkey Anti-Rabbit IgG (H+L) | Cat#711-175-152, Jacksonimmun | | 1:500 |
| TRITC AffiniPure Donkey Anti-Goat IgG (H+L) | Cat#705-025-147, Jacksonimmun | | 1:500 |

| | | | |
|---|---|---|---|
| Alexa Fluor$^{TM}$ 488 Donkey anti-Goat IgG（H+L） | Cat#A11055, Thermo Fisher Scientific | | 1:500 |
| Alexa Fluor$^{TM}$ 488 Goat anti-Rabbit IgG（H+L） | Cat#A11034, Thermo Fisher Scientific | | 1:500 |
| DAPI | Cat#D9542, Sigma | | 1:500 |

**Software and Algorithms**

| Software name | Developer | Source |
|---|---|---|
| ImageJ | NIH | https://imagej.en.softonic.com/ |
| GraphPad Prism | GraphPad Software | https://www.graphpad.com/ |
| Adobe Photoshop | Adobe | https://www.adobe.com/ |
| Adobe illustrator | Adobe | https://www.adobe.com/ |
| IBM SPSS | IBM | https://www.ibm.com/cn-zh/spss |
| SnapGene | SnapGene | https://www.snapgene.com/ |
| ZEN 2012 | ZEISS | https://www.zeiss.com.cn/ |
| BD FACSDiva | BD Biosciences | https://www.bdbiosciences.com/ |
| NIS-Elements Viewer | Nikon | https://www.nikon.com/ |
